# PIN1 isomerisation of BRCA1 promotes replication fork protection

**DOI:** 10.1101/478511

**Authors:** Manuel Daza-Martin, Mohammed Jamshad, Katarzyna Starowicz, Anoop Singh Chauhan, James F.J. Beesley, Jennifer L. Coles, Grant S. Stewart, Ruth M. Densham, Joanna R. Morris

## Abstract

BRCA1, BRCA2 and a subset of Fanconi’s Anaemia proteins act to promote RAD51-mediated protection of newly synthesised DNA at stalled replication forks from degradation by nucleases. How BRCA1 contributes, how it is regulated and whether replication fork protection relates to, or differs from, the roles BRCA1 has in homologous recombination is not clear. Here we show that the canonical BRCA1-PALB2 interaction is not required for fork protection and instead we demonstrate that the ability of BRCA1 to protect nascent DNA is regulated in an unexpected fashion through conformational change mediated by the phosphorylation-directed prolyl isomerase, PIN1. BRCA1 isomerisation enhances BRCA1-BARD1 interaction with RAD51 and consequently RAD51 presence at stalled replication structures. Our data suggest BRCA1-BARD1 promotes fork protection in part by enhancing the RAD51 synapse. Patient missense variants in the regulated BRCA1-BARD1 regions similarly show poor nascent strand protection but proficient homologous recombination, defining novel domains required for fork protection in BRCA1-BARD1 associated with cancer predisposition. Together these findings reveal a previously unrecognised pathway that governs BRCA1-mediated replication fork protection.

## Introduction

Genomic integrity is constantly under threat by problems that the replication fork might encounter^1^. Fork progression can be slowed by conflicts with transcription, deoxyribonucleotide (dNTP) shortage or difficult to replicate sequences, frequently causing fork stalling. To avoid replication stress, a set of responses act to prevent replication forks from collapsing, amongst which is fork remodelling and subsequent nascent strand protection.

In remodelling newly made DNA strands anneal and a four-way, chicken foot structure is evident in electron microscopy (also called nascent strand exchange or fork reversal ^2,3^). Agents that cause replicative stress or compromise Pol alpha function result in a proportion of forks reversing ^2-5^, reviewed in ^6,7^. The regressed arm of nascent DNA in reversed forks resembles a single-ended DNA double strand break, and while some resection of the structure is required for replication fork restart, the reversed fork is protected from excessive resection by RAD51. Several factors contribute to RAD51-mediated fork protection including BRCA1/2, FANCA/D2, RAD51 paralogs, BOD1L, SETD1A WRNIP and Abro1^8-14^.

An emerging theme is that stability of the RAD51-nucleofilament is critical to the protection of stalled forks ^8,15^. Increased dissociation of RAD51 mutants compromises their ability to prevent fork degradation ^16-18^. Also, factors such as BOD1L stabilise RAD51 on ssDNA and promote fork protection^19^, while others, such as RADX, compete with or dissociate RAD51, so that their depletion rescues fork protection of BRCA-defective cells ^20,21^.

Failure of replication fork protection is associated with genome instability, but whether it constitutes a distinct tumour suppressor pathway is not yet clear. Restoration of replication fork protection in BRCA-deficient cells is linked to chemotherapy resistance in some cell types and contexts ^22-24^. Thus further mechanistic understanding of fork protection is thus needed to inform cancer patient care.

BRCA1 is found at ongoing and stalled replication forks ^25-27^ and like BRCA2, contributes to RAD51-mediated defence of stalled forks from MRE11-dependent degradation^9^. Cells with BRCA1 haplo-insufficiency show more frequently degraded forks ^28^, suggesting that poor fork degradation may allow genome instability in the normal and pre-cancerous cells of *BRCA1* mutation carriers. However, how BRCA1 contributes to fork protection is not clear.

The data presented here reveal BRCA1 promotes the protection of nascent DNA at stalled replication forks independently of the canonical BRCA1-PALB2 interaction. Instead we find that an enhanced direct interaction between BRCA1-BARD1 and RAD51 is required for fork protection and that this enhanced interaction is dependent on conformational changes catalysed by the phosphorylation directed prolyl isomerase, PIN1. These newly identified BRCA1 and BARD1 regions required for fork protection harbour patient variants associated with familial and sporadic cancer which can inhibit fork protection. Additionally our data infers that the enhanced RAD51-BARD1-BRCA1 interaction stabilises RAD51-synapse-like structures at stalled forks.

## Results

### The BRCA1-BARD1 heterodimer promotes fork protection through the RAD51 binding region and not the PALB2-BRCA2 interaction region

During homologous recombination (HR) of DNA double-strand breaks (dsDNA breaks), BRCA1-BARD1 regulates RAD51 localisation and loading through PALB2-BRCA2 ^29-31^ and also directly interacts with RAD51 to promote invasion and D-loop formation ^32^. BRCA1, PALB2 and BRCA2 are found at stalled and collapsed replication forks ^33^ and while the PALB2-BRCA2 interaction is required for fork protection^34^, it is unclear whether the BRCA1-PALB2 interaction is relevant. To address this question we examined nascent DNA at hydroxyurea (HU) stalled forks. Newly synthesized DNA was labelled using CldU before stalling with 5 mM HU for 3 hours. After DNA fibre-spreading, the length of the label was measured as an indicator of the stability of newly synthesized DNA at stalled replication forks ^8^ ^9^ (Fig 1A). Replication stalling caused a dramatic shortening of CldU tract lengths in BRCA1 or PALB2 depleted cells (just 8.6 and 8.7 μm respectively in PALB2 and BRCA1 depleted cells compared to 13.7 μm in controls) (Fig 1B-C). BRCA1 and PALB2 directly interact through coiled-coil motifs ^30,31^. The BRCA1-M1141T point mutation prevents the PALB2 interaction, fails to support HR-repair ^31^, and shows increased sensitivity to the inter-strand cross-linker Cisplatin in colony survival assays (Supplementary Fig 1A-C). Similarly, an N-terminal coiled-coil deletion mutant of PALB2 (ΔNT-PALB2) fails to interact with BRCA1 ^30,31^ and shows increased sensitivity to Cisplatin in colony survival assays (Supplementary Fig 1D). Surprisingly, when we used these interaction mutants to assess fork protection using the CldU incorporation assay, both BRCA1-M114T and ΔNT-PALB2 were able to stabilise stalled replication fork tract lengths in BRCA1 or PALB2 siRNA-depleted cells to similar lengths as controls (Fig 1D-G). In contrast to this, the BRCA1-PALB2 canonical interaction is required for replication after HU incubation as both the M1411T-BRCA1 and ΔNT-PALB2 mutations exhibit a reduced proportion of replication fork restart and increased fork stalling (Supplementary Fig 1E-H). Therefore, while both BRCA1 and PALB2 are required to protect nascent DNA at stalled replications forks, fork restart, but not fork protection, is dependent on the canonical interaction between BRCA1 and PALB2.

**Figure 1.**
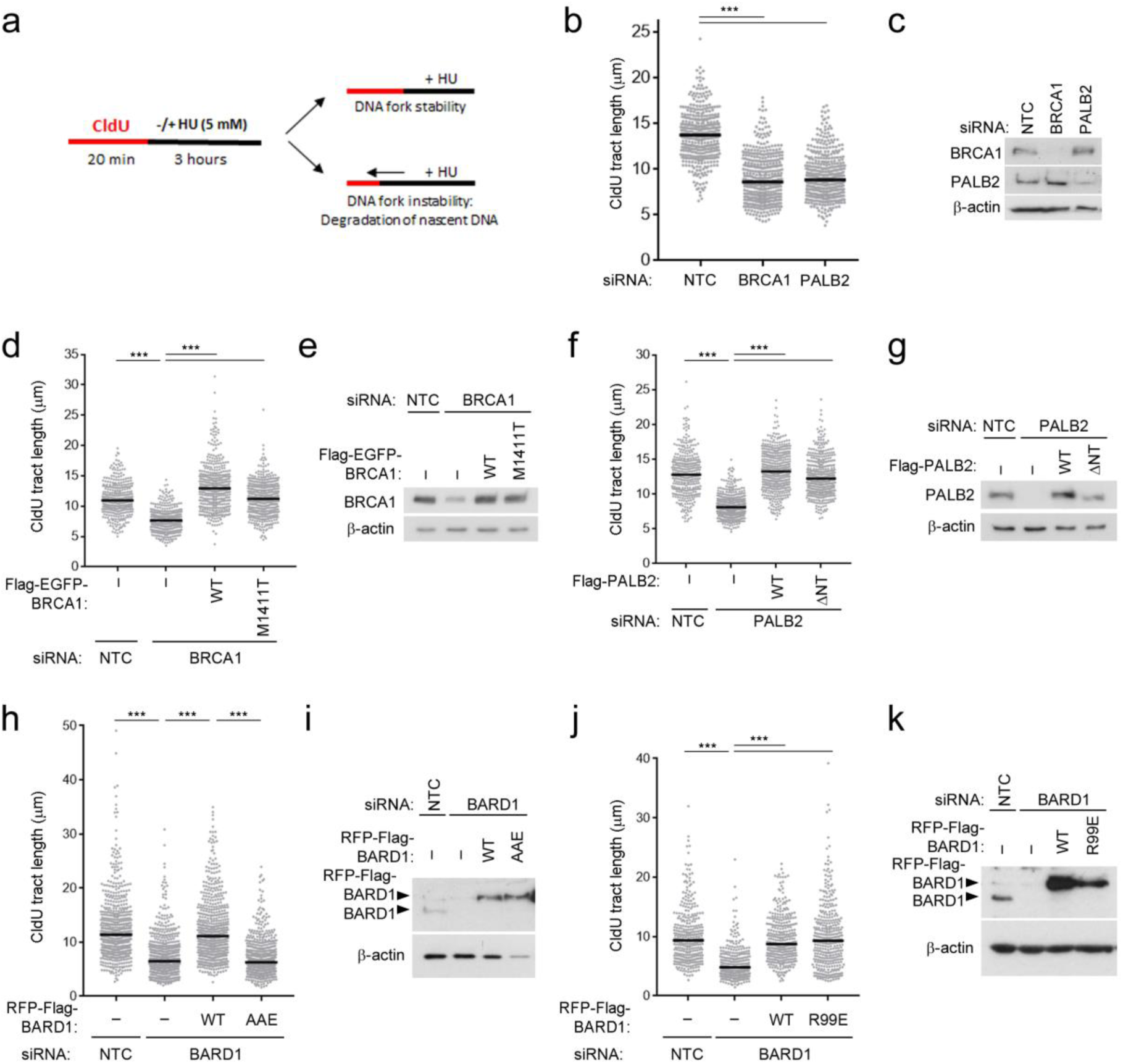
The BRCA1-BARD1 heterodimer promotes fork protection through the RAD51 binding region and not the PALB2-BRCA2 interaction region. a. Diagram to illustrate the replication fork protection assay by assessing CldU tract lengths following exposure to HU. b. CldU fibre tract lengths were measured from U20S cells depleted for BRCA1 or PALB2 treated with 5 mM HU for 3 hours. N=430 fibres from 3 independent experiments, bars indicate median. c. Western blot to show BRCA1 and PALB2 depletions for B d) CldU fibre tract lengths were measured from U20S cells depleted for BRCA1 and complemented with WT or M1411T-Flag-EGFP-BRCA1 and treated with 5 mM HU for 3 hours. N=340 fibres from 3 independent experiments, bars indicate median. e. Western blot to show BRCA1 depletions and complementation for D. f. CldU fibre tract lengths were measured from U20S cells depleted for PALB2 and complemented with WT or ΔNT-Flag-PALB2 and treated with 5 mM HU for 3 hours. N=360 fibres from 3 independent experiments, bars indicate median. g. Western blot to show PALB2 depletions and complementation for F. h. CldU fibre tract lengths were measured from U20S cells depleted for BARD1 and complemented with WT or RAD51-binding deficient AAE-RFP-Flag-BARD1 and treated with 5 mM HU for 3 hours. N=600 fibres from 3 independent experiments, error bars = median. i. Western blot to show BARD1 depletions and complementation for H. j. CldU fibre tract lengths were measured from U20S cells depleted for BARD1 and complemented with WT or R99E-RFP-Flag-BARD1 and treated with 5 mM HU for 3 hours. N=350 fibres from 3 independent experiments, bars indicate median. k. Western blot to show BARD1 depletions and complementation for J.

Both BRCA1 and BARD1 carry regions able to directly interact with RAD51 ^32,35^. BARD1 residues F133, D135 and A136 form part of an interaction face with RAD51, and are required for D-loop formation in HR and for mitomycin C, Olaparib and HU resistance ^32^ (Supplementary Fig 1I-J). We assessed cells complemented with BARD1 bearing substitutions in this RAD51-binding site (F133A-D135A-A136E, termed AAE-BARD1). Depletion of BARD1 caused a similar decrease in CldU tract lengths as that seen following BRCA1 depletion (11.4 µm in controls vs 6.5 µm in siBARD1. after HU treatment and complementation with WT-BARD1 but not AAE-BARD1 supported protection of nascent strands (11.1 µm vs 6.3 µm) (Fig 1H-I). In contrast when we assessed the requirement for the BRCA1-BARD1 E3 ubiquitin ligase function we found that complementation with R99E-BARD1, a mutant that specifically disrupts E3 ligase activity without disrupting the BRCA1-BARD1 heterodimer ^36^, was sufficient to restore longer CldU tract lengths following HU-induced fork stalling (9.6 µm Vs 8.7 µm in cells with WT-BARD1) (Fig 1J-K). Therefore, protection of stalled replication forks requires a functional RAD51 binding site in BARD1 but not the BRCA1-BARD1 E3 ubiquitin ligase activity, nor the canonical BRCA1-PALB2 interaction.

### The BRCA1 serine 114 phosphorylation site is required for the protection of nascent DNA

BRCA1 and BARD1 are phosphorylated at residues that are structurally potentially close to the BARD1-RAD51 interaction site: at S148 in BARD1 ^37^ and S114 in BRCA1 ^38,39^. We tested whether these sites are required in fork protection by mutation and found substitution to alanines (S114A-BRCA1 or S148A-BARD1) shortened CldU tract lengths following HU-treatment but not in the absence of HU-treatment (Fig 2A-B and Supplementary Fig 2A-C). In contrast, mutation to aspartate (D) as a phospho-mimic was able to support fork protection in S114D-BRCA1 complemented cells but not in S148D-BARD1 complemented cells (Fig 2C-D and Supplementary Fig 2D-E), supporting a role for BRCA1 phosphorylation in fork protection.

**Figure 2.**
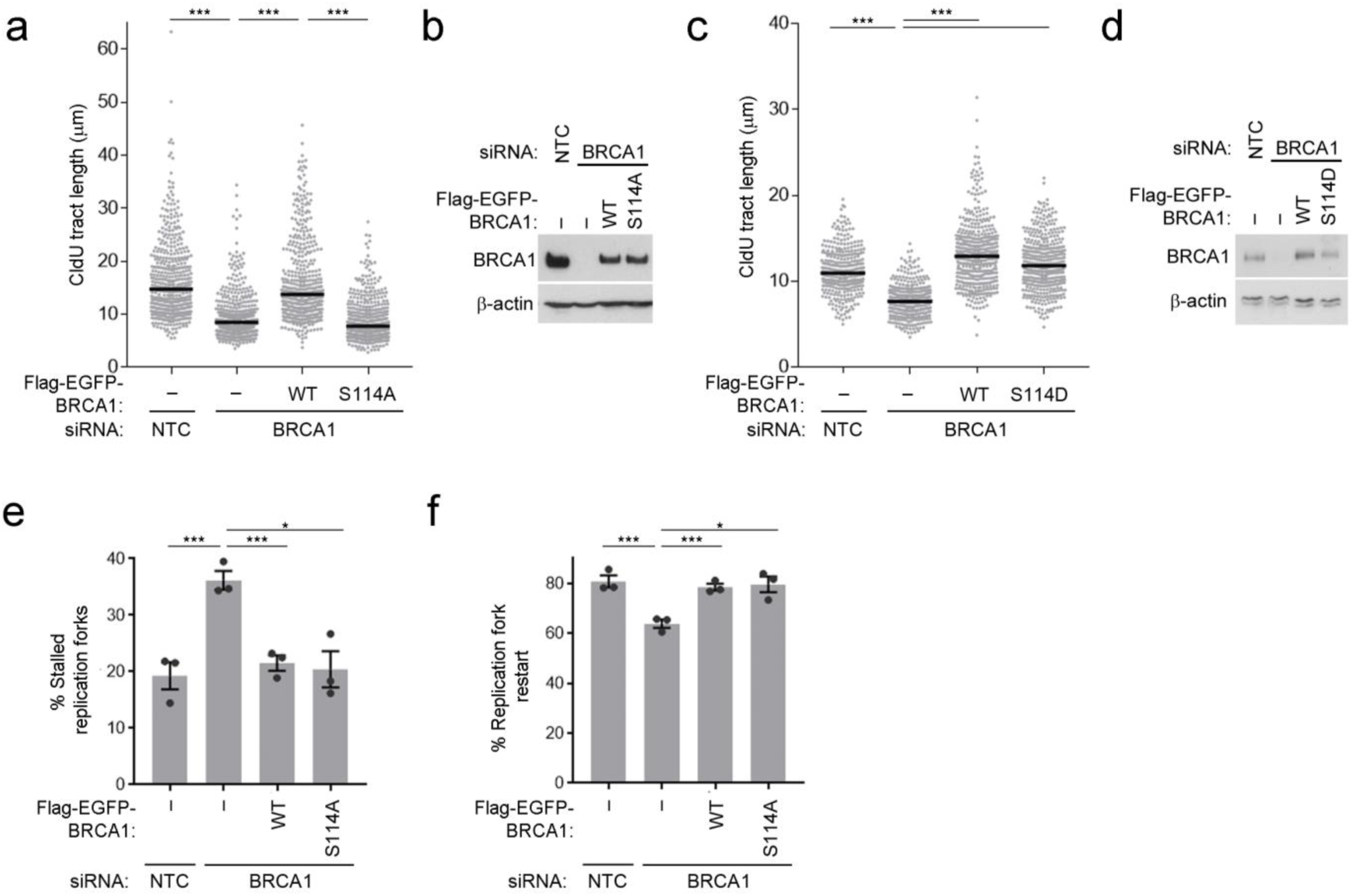
The BRCA1 serine 114 phosphorylation site is required for the protection of nascent DNA. a) CldU fibre tract lengths were measured from U20S cells depleted for BRCA1 and complemented with WT or S114A-Flag-EGFP-BRCA1 treated with 5 mM HU for 3 hours. N=460 fibres from 3 independent experiments, bars indicate median. b. Western blot to show BRCA1 depletions and complementation for A. c. CldU fibre tract lengths were measured from U20S cells depleted for BRCA1 and complemented with WT or S114D-Flag-EGFP-BRCA1 treated with 5 mM HU for 3 hours. N=340 fibres from 3 independent experiments, bars indicate median. d. Western blot to show BRCA1 depletions and complementation for C. e. The % stalled replication forks were measured from 3-independent experiments in U20S cells depleted for BRCA1 and complemented with WT or S114A-Flag-EGFP-BRCA1. Grey bars indicate mean, error bars are SEM. f. The % replication forks able to restart after release from 3 hr of 5 mM HU were measured from 3-independent experiments in U20S cells depleted for BRCA1 and complemented with WT or S114A-Flag-EGFP-BRCA1. N=3. Grey bars indicate mean, error bars are SEM.

The S114A-BRCA1 and WT-BRCA1 had similar levels of recruitment to sites of active replication identified by the incorporation of the nucleotide analogue CldU following a pulse treatment (Supplementary Fig. 2F) and both mutant and WT-BRCA1 interacted with BARD1 to the same degree (Supplementary Fig 2G).

BRCA1 has several roles at replication forks ^40^ and depletion of BRCA1 increased the stalling of ongoing replication and also reduced replication restart following release from short term (3 hour) exposure to 5 mM HU. Complementation with either WT-BRCA1 or S114A-BRCA1 restored both ongoing forks and fork restart (Fig 2E-F) suggesting that the S114-site is not significant to these aspects of replication stress. Fork protection defects can be restored in BRCA1/2 deficient cells by inhibition of MRE11 3’-5’ nuclease activity with Mirin ^9^, and similarly Mirin restored long tract lengths in S114A-BRCA1 complemented cells (from 7.6 μm to 11.3 μm) (Supplementary Fig 2H-1), implicating the S114-site in the protection of nascent DNA from nuclease activity.

### Phosphorylation of BRCA1 at serine 114 promotes PIN1 interaction

We generated an antibody to a phosphorylated-S114 BRCA1 peptide (p-S114) which was able to detect immunoprecipitated WT-BRCA1 and not S114A-BRCA1 (Fig 3A), confirming phosphorylation at S114. Intriguingly, the BRCA1 phospho-S114 site lies within an S-P motif which is a minimal consensus site for the phospho-peptidyl-prolyl isomerase (PPIase), PIN1, and BRCA1 and BARD1 have previously been enriched from lysates using recombinant PIN1 ^41^. We were able to confirm their interaction by immunoprecipitation of the BARD1-BRCA1-PIN1 complex from cells over-expressing RFP-Flag-BARD1 (Fig 3B). Since protein interactions with the full length PIN1 enzyme are transient, we generated a GST-fusion of the PIN1 WW phospho-binding domain (GST-WW) without the PPIase domain, and a corresponding GST-W34A mutant form which is deficient in phospho-binding, to explore PIN1 interactions further ^42^ (Supplementary Fig 3A). Purification of WT-BRCA1 and the S114D-BRCA1 phospho-mimic, but not S114A-BRCA1 was achieved by GST-WW, but not GST-W34A (Fig. 3C and Supplementary Fig 3B). Furthermore, the interaction between either exogenous Flag-EGFP-BRCA1 or endogenous BRCA1 and recombinant PIN1-WW domain was increased following HU treatment (Fig 3D and Supplementary Fig 3C). The interaction of endogenous PIN1 isomerisation of BRCA1 promotes replication fork protection.

**Figure 3.**
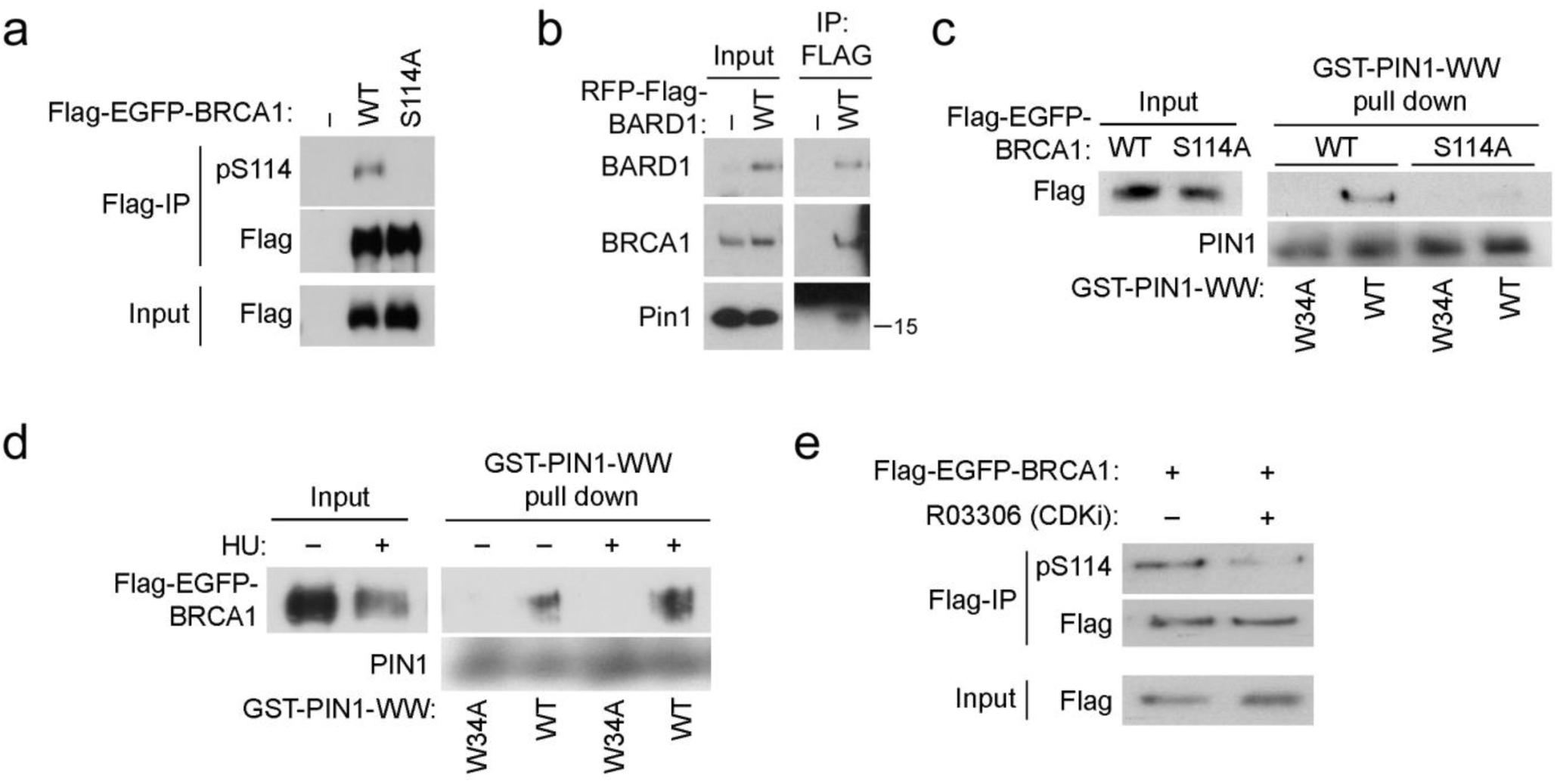
Phosphorylation of BRCA1 at serine 114 promotes PIN1 interaction. a. BRCA1 phosphorylation at S114 was measured in HEK293 cells expressing Flag-EGFP-BRCA1. FLAG-immunoprecipitation of WT or S114A Flag-EGFP-BRCA1 was probed for pS114-BRCA1 by western blot. b. FLAG-immunoprecipitations from U20S cells expressing RFP-Flag-BARD1 demonstrated co-immunoprecipitation of endogenous BRCA1 and PIN1. c. Glutathione-Sepharose beads bound with the GST-fused-WW domain of PIN1 were used to pull-down WT, but not S114A Flag-EGFP-BRCA1, from U20S cell lysates. Beads bound by GST-W34A WW-domain were used as a negative control. d. Glutathione-Sepharose beads bound with the GST-fused-WW domain of PIN1 were used to pull-down WT from HEK293 cell lysates treated with or without 3 mM HU for 6 hours. Beads bound by GST-W34A WW-domain were used as a negative control. e. BRCA1 phosphorylation at S114 was measured in HEK293 cells expressing Flag-EGFP-BRCA1 and treated with 5 μM CDK1/2 inhibitor RO-3306 for 30 minutes. FLAG-immunoprecipitation of WT Flag-EGFP-BRCA1 was probed for pS114-BRCA1 by western blot.

BRCA1 with PIN1-WW domain was lost following treatment of the cell lysate with phosphatase (Supplementary Fig 3C, final lane). In contrast, while WT-BARD1 was also purified by GST-WW and not GST-W34A, this interaction was not lost on mutation of the RAD51 proximal phosphorylation site S148 (S148A-BARD1) (Supplementary Fig 3D) suggesting that pS148-BARD1 is not a PIN1 interaction site. Together these data suggest that BRCA1 is phosphorylated at S114 in response to HU and that this phosphorylation promotes the ability of the PIN1-WW domain to purify BRCA1.

The BRCA1 S114 site lies within a loose CDK1/2 consensus site (S-P-x-x-x-K) and treatment of cells with the CDK1/2 inhibitor (Roscovitine), the CDK1 inhibitor (RO-3306) or with CDK1 siRNA reduced the affinity of the p-S114 antibody for immunoprecipitated BRCA1 and the ability of the GST-WW domain of PIN1 to purify BRCA1 from cell lysates (Fig 3E and Supplementary Fig 3E-F). Similarly incubation of recombinant CDK2/Cyclin A with recombinant WT or S114A His-BRCA1_1-300_-BARD1_26-142_specifically phosphorylated WT-BRCA1 but not S114A-BRCA1 (Supplementary Fig 3G). These data indicate CDK-mediated phosphorylation at S114 contributes to the ability of PIN1 to interact with BRCA1.

### PIN1 regulates the BRCA1-BARD1 heterodimer to promote fork protection

PIN1 is expressed at low levels in non-proliferating cells and increases with proliferative capacity (reviewed in^43^) and studies that have isolated proteins on nascent DNA (iPOND) have shown PIN1 enrichment following treatment with HU ^20,25,44^. Inhibition of PIN1 with Juglone, or depletion of PIN1 by siRNA, led to a shortening of CldU tracts following HU-treatment consistent with a fork protection defect. This was epistatic with BRCA1 or BARD1 loss (Fig 4A-B and Supplementary Fig 4A-C) suggesting that PIN1’s role in fork protection lies in the same pathway as BRCA1-BARD1. While a common outcome of PIN1 interaction is altered protein stability ^41,45-48^, we saw no gross impact on BRCA1 or BARD1 levels following PIN1 depletion (Fig 4B and Supplementary Fig 4C).

**Figure 4.**
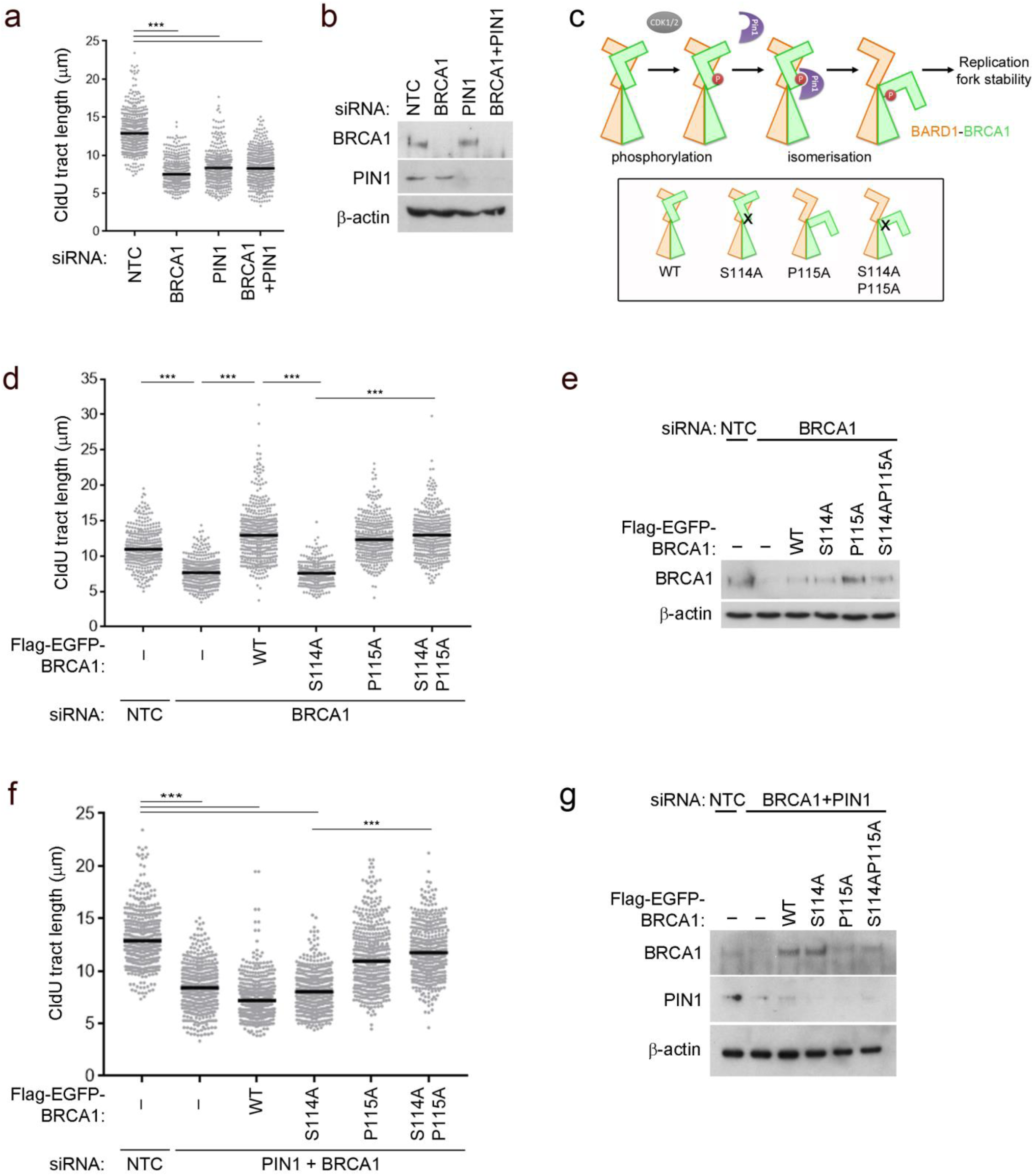
PIN1 regulates the BRCA1-BARD1 heterodimer to promote fork protection. a. CldU fibre tract lengths were measured from U20S cells depleted for BRCA1 and/or PIN1 and treated with 5 mM HU for 3 hours. N=400 fibres from 3 independent experiments, bars indicate median. b. Western blot to show BRCA1 and PIN1 depletions described for A. c. Schematic to illustrate CDK1/2 (grey) phosphorylation at S114 (red) and subsequent PIN1 (purple) isomerisation events on the BRCA1 (orange) and BARD1 (green) N-termini. Boxed cartoons illustrate the phosphorylation and isomerisation mutants of BRCA1. d. CldU fibre tract lengths were measured from U20S cells depleted for BRCA1 and complemented with Flag-EGFP-BRCA1 variants as indicated and treated with 5 mM HU for 3 hours. N=>250 fibres from 3 independent experiments, bars indicate median. e. Western blot to show BRCA1 depletions and complementation for D. f. CldU fibre tract lengths were measured from U20S cells co-depleted for BRCA1 and PIN1, and complemented with Flag-EGFP-BRCA1 variants as indicated. Cells were treated with 5 mM HU for 3 hours. N=450 fibres from 3 independent experiments, bars indicate median. g. Western blot to show BRCA1 and PIN1 depletions and complementation as described in F.

In folded proteins, peptide bonds preceding residues other than proline (non-prolyl bonds) overwhelmingly favour the *trans* form, and *cis* bonds are rare in folded proteins ^49^. In contrast, due to the physical constraints of proline’s unique 5-membered ring on the peptide backbone, peptide bonds preceding proline (prolyl bonds) more often adopt the *cis* conformation ^50^. PIN1 is the only phospho-targeted PPIase; it specifically recognises phospho-S/T-P motifs ^51^ before catalysing a *cis-trans* conformational change on the peptidyl-prolyl bond in the peptide backbone _52-57_. An experimental approach often used to examine potential requirements for the *trans*-isomer is to substitute the proline of the target with other amino acids ^58,59^. We tested the requirement for proline isomerisation to *trans* by mutating BRCA1 P115 to an alanine to constitutively present a *trans* backbone prior to residue 115 (Fig 4C).

P115A-BRCA1 was able to complement BRCA1 depletion preventing shortening of nascent tracts after HU exposure, suggesting no negative impact of *trans*-isomerisation or proline loss at this position (Fig 4D-E). Importantly inclusion of P115A with the S114A mutation resulted in normal CldU tract lengths in BRCA1-depleted cells (Fig 4D), showing that inclusion of an alanine at P115 can overcome the requirement for serine at 114. These data suggest *trans* isomerisation of P115 bypasses the need for S114 phosphorylation. In addition, mutation of P115 to another amino acid, cysteine, also overcame the requirement for the S114-site in fork protection (Supplementary Fig 4D-E) supporting the hypothesis that this is due to presenting BRCA1 in a *trans* conformation at position 115 rather than due to a specific effect of alanine at P115.

We next addressed the degree to which the failure to maintain nascent DNA in cells with repressed PIN1 is due to isomerisation at P115-BRCA1. In BRCA1 and PIN1 co-depleted cells expression of P115A or S114A-P115A-BRCA1 resulted in lengthened average tract lengths (11.0 μm in P115A and 11.7μm in S114A-P115A cells) compared to cells with BRCA1+PIN1 co-depletion (8.3 μm). However, lengths were not fully restored to that of nascent strands in control cells (12.9 μm) (Fig. 4F-G). Thus the requirement for PIN1 in replication fork protection can be largely overcome by expression of a *trans*-locked mutant of BRCA1 at position P115, suggesting the majority of the need for the PIN1 PPIase is through BRCA1.

### BRCA1-BARD1 isomerisation enhances direct RAD51 binding and promotes accumulation at nascent DNA

Using recombinant purified His-BRCA1_1-500_-BARD1_27-327_ (Supplementary Fig 5A) we used limited trypsin proteolysis to assess the *cis-trans* conformational change predicted by mutation of P115 to alanine. The P115A version of BRCA1 was more resistant to trypsin, transitioned more slowly through the ~18 kDa intermediate at 3-15 minutes post addition of trypsin, and formed the ~15 kDa fragment maximally at 30 min post-trypsin compared to 3 minutes with WT-BRCA1. The same size kDa fragments are formed by both WT and P115A suggesting that there is no gross change of domain structure but rather a change in accessibility to the same trypsin digestion sites (Fig 5A). These data are consistent with a subtle structural conformational difference between WT and P115A-BRCA1. Given the potential proximity of P115 in BRCA1 to the RAD51 binding site of BARD1 we next asked whether *cis-trans* isomerisation of BRCA1 would affect RAD51 binding to the BRCA1-BARD1 heterodimer. WT and P115A recombinant His-BRCA1_1-500_-BARD1_27-327_ were incubated with active recombinant RAD51 and the direct protein-protein interaction assessed by His-pull down. Our analysis, demonstrated greater recombinant RAD51 binding to the *trans* P115A BRCA1-BARD1 heterodimer compared to the WT-BRCA1-BARD1 heterodimer (Fig 5B).

**Figure 5.**
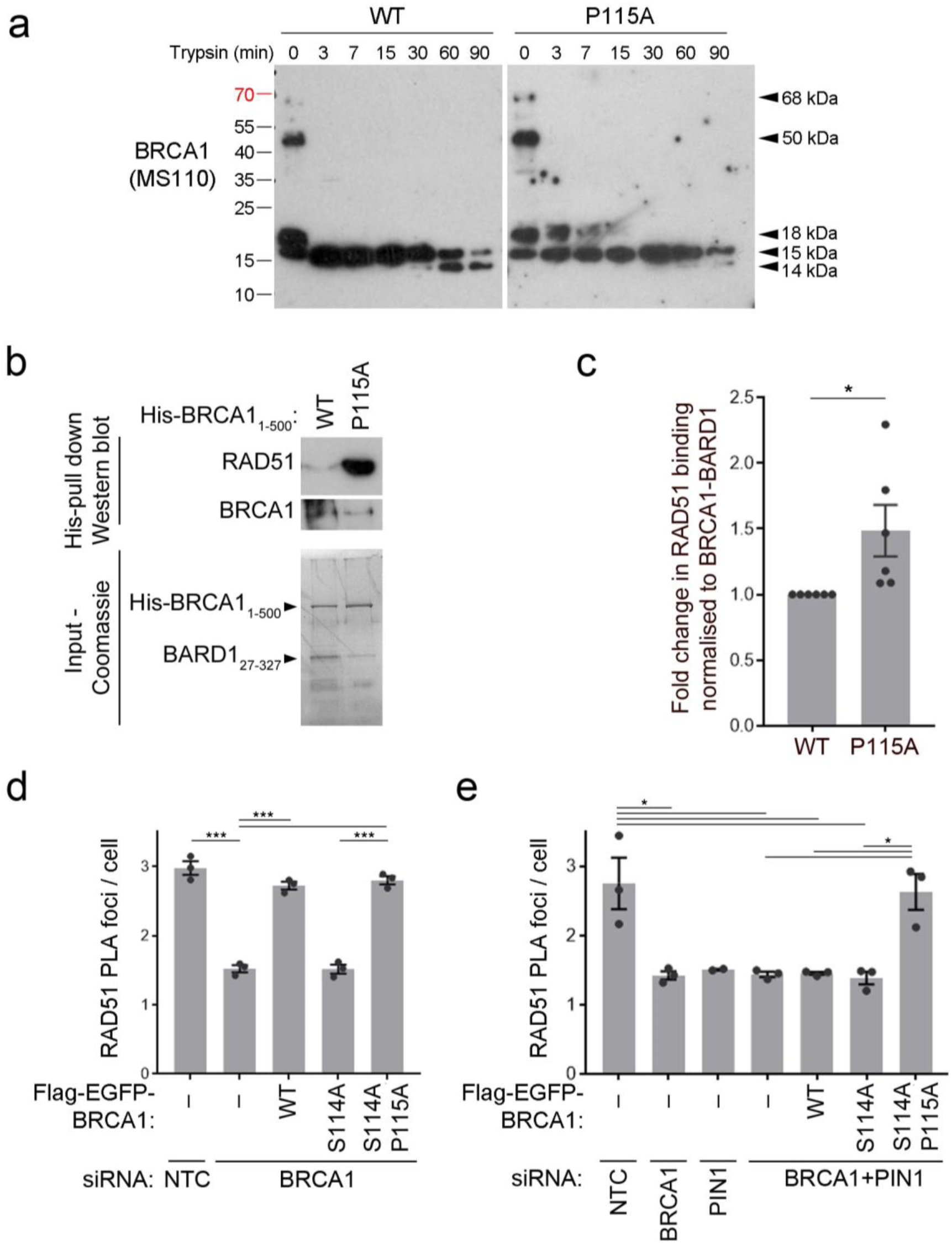
BRCA1-BARD1 isomerisation enhances direct RAD51 binding and promotes accumulation at nascent DNA. a. Recombinant WT or P115A His-BRCA11-500 and BARD127-327 were incubated with Trypsin and samples taken at the times indicated. The limited proteolysis profiles for WT v P115A BRCA1 were assessed by western blot using monoclonal BRCA1 MS110 antibody. b. Recombinant WT or P115A His-BRCA11-500 and BARD127-327 were incubated with recombinant active RAD51. The ability of BRCA1-BARD1 to bind RAD51 was assessed by His-purification of the BRCA1-BARD1-RAD51 complex, followed by western blot as indicated c. Quantification of Flag-Immunoprecipitation of RAD51 bound to WT or P115A Flag-EGFP-BRCA1 from HEK293 cells. N=6. Grey bars indicate mean, error bars are SEM. d. RAD51 colocalisation with nascent DNA, marked by pulse labelling with EdU, was measured using the proximity ligation assay (PLA) in U20S cells treated with 5 mM HU for 4 hours, depleted for BRCA1 and complemented with Flag-EGFP-BRCA1 variants as indicated. Mean number of RAD51/EdU-Biotin PLA foci per cell was measured from 3-independent assays. Grey bars indicate mean. Error bars are SEM. e. RAD51 colocalisation with nascent DNA, marked by pulse labelling with EdU, was measured using the proximity ligation assay (PLA) in U20S cells treated with 5 mM HU for 4 hours, depleted for BRCA1 and/or PIN1, and complemented with Flag-EGFP-BRCA1 variants as indicated. Mean number of RAD51/EdU-Biotin PLA foci per cell was measured from 3-independent assays. Grey bars indicate mean. Error bars are SEM.

We then wanted to address the BRCA1-BARD1-RAD51 interaction in cell lysates. Immunoprecipitation of P115A-BRCA1 in complex with WT-BARD1 showed a 1.5-fold enhancement in the interaction with RAD51 compared to WT-BRCA1 (Fig 5C and Supplementary Fig 5B). To assess any influence on RAD51 at nascent DNA following HU-treatment we used proximity linked ligation assay (PLA) with antibodies to RAD51 and Biotin conjugated to EdU ^60^. This showed that BRCA1-depleted cells complemented with S114A-P115A-BRCA1 showed RAD51 PLA foci levels similar to WT complementation and greater than the S114A-BRCA1 complementation or BRCA1 depleted cells (Fig 5D and Supplementary Fig 5C). Thus a *trans*-locked form of BRCA1 at the peptidyl-115 bond overcomes the requirement for a functional phosphorylation site at S114 in promoting RAD51 accumulation at nascent DNA. Likewise in cells depleted for BRCA1 and PIN1 the RAD51 PLA foci levels were restored by complementation of S114A-P115A-but not S114A-BRCA1 (Fig 5E). These data indicate that isomerisation of BRCA1 contributes to the presence of RAD51 at stalled forks.

### Loss of BRCA1-isomerisation leads to genomic instability and increased sensitivity to replication stress agents

Mutation of the RAD51 binding site in BARD1 (AAE) leads to increased sensitivity to both replication stress (HU) and DNA damaging agents such as Olaparib and Camptothecin that require HR-directed repair ^32^ (Supplementary Fig 1I&J). When we assessed the requirement of the S114-site in BRCA1 for cell survival in response to PARP inhibitors (Olaparib, Veliparib or 4AN) or replication stress agents (HU or Aphidicolin) we found complemented cells were resistant to PARP inhibitors and formed RAD51 foci after Olaparib treatment but were sensitive to overnight treatment with replication stress-inducing agents (Supplementary Fig 6A-C, Fig 6A-B). Since prolonged treatment with HU causes replication fork collapse and the formation of double strand breaks, we also assessed colony survival in response to conditions that promote fork stalling but not collapse (3 hour treatment of 5 mM HU)^61^, conditions identical to those used in our fork protection assays. Intriguingly, even in asynchronous cells treated with short-term HU, the S114A-BRCA1 complemented cells showed reduced colony formation (Fig 6D). Moreover inclusion of the P115A mutation on the S114A background was sufficient to rescue the S114A defect in survival seen in response to both 3 hour and overnight treatment with HU (Fig 6D-F). In previous reports low-dose HU experiments have had little-impact on cell survival (for example of BRCA2-deficeint cells^9^). We have discounted several replication defects (restart, fork speed, stalling) and in our hands resistance to both short and long-term HU exposure closely correlates with fork protection.

**Figure 6.**
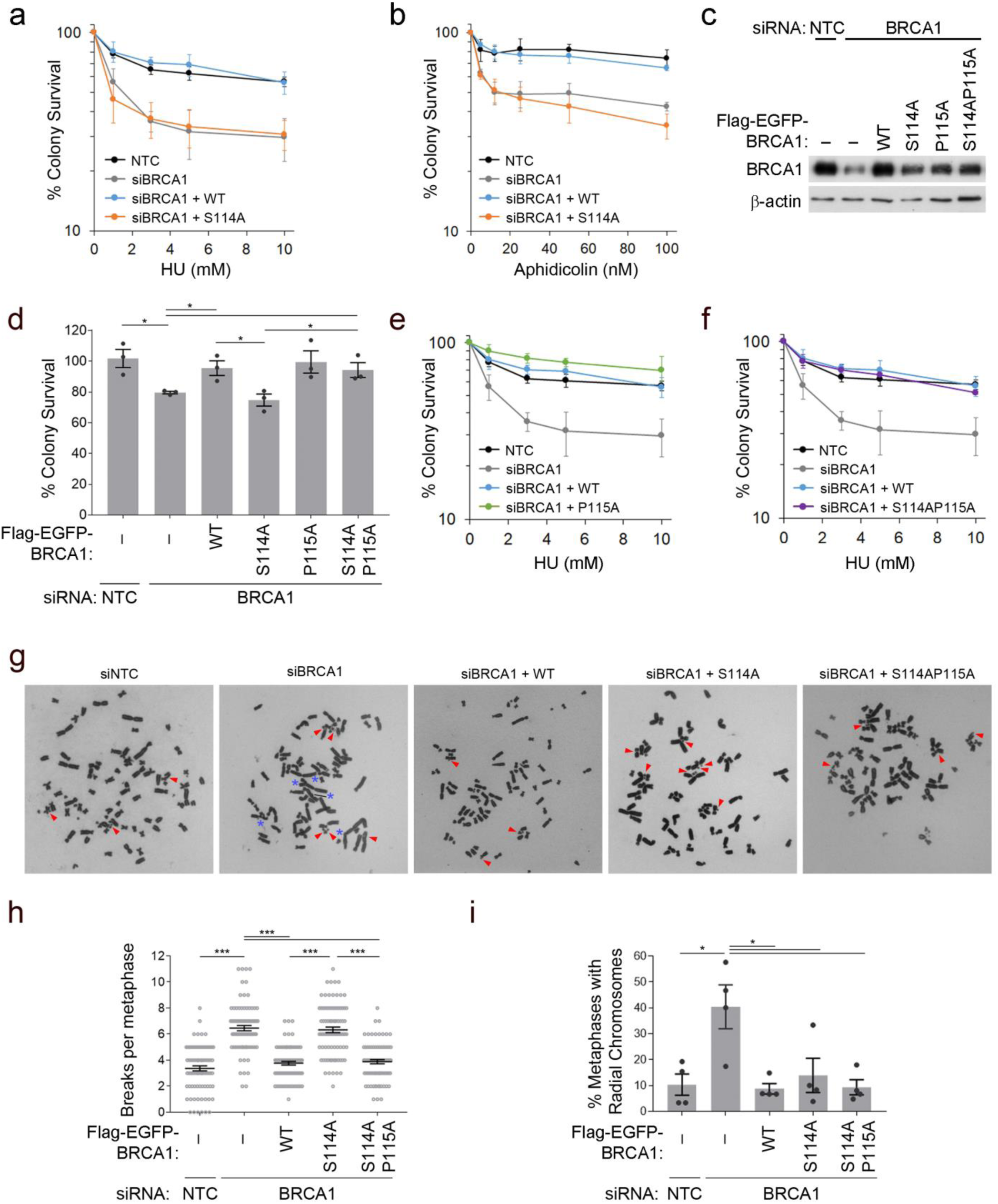
Loss of BRCA1-isomerisation leads to genomic instability and increased sensitivity to replication stress agents. a. Colony survival following 16 hour treatment with HU was measured in HeLa cells depleted for BRCA1 and complemented with WT or S114A Flag-EGFP-BRCA1. N=4, error bars are SEM. b. Colony survival following 16 hour treatment with Aphidicolin was measured in HeLa cells depleted for BRCA1 and complemented with WT or S114A Flag-EGFP-BRCA1. N=4, error bars are SEM. c. Western blot to show BRCA1 depletions and complementation for A-B. d. Colony survival following 3 hour treatment with 5mM HU was measured in U20S cells depleted for BRCA1 and complemented with WT or Flag-EGFP-BRCA1 variants as indicated. N=3, error bars are SEM. e. Colony survival following 16 hour treatment with HU was measured in HeLa cells depleted for BRCA1 and complemented with WT or P115A Flag-EGFP-BRCA1. N=3, error bars are SEM. f. Colony survival following 16 hour treatment with HU was measured in HeLa cells depleted for BRCA1 and complemented with WT or S114AP115A Flag-EGFP-BRCA1. N=3, error bars are SEM. g. Representative images of metaphase spreads from U20S cells depleted for BRCA1 and complemented with WT or S114A or S114AP115A Flag-EGFP-BRCA1. Cells were treated with 5 mM HU for 4 hours before being blocked with Colcemid to trap cells in metaphase. Red arrows indicate chromosome and chromatid breaks. Blue asterisks mark radials. h. Metaphase spreads from G were scored for the number of chromosome or chromatid breaks per metaphase. 80 metaphases from 3-independent experiments were scored. Bars indicated mean and error bars are SEM. i. Metaphase spreads described in G were scored for the % of metaphases showing 1 or more radial chromosomes. Data is from 4-independent experiments. Grey bars indicate mean and error bars are SEM.

These data lead us to investigate the role of the phosphorylation-isomerisation region of BRCA1 in genome stability. As expected depletion of BRCA1 increased both the average number of breaks per metaphase (from 3.4 in controls to 6.5 in BRCA1-depleted cells) and the percentage of metaphases with chromatid exchanges (10.3 % in controls to 40.4 % in siBRCA1) (Fig 6G-I). Strikingly, cells complemented with WT-BRCA1 or S114A-P115A-BRCA1 double mutant, but not S114A-BRCA1, were able to restore the average number of breaks per metaphase to levels seen in controls (Fig 6G-H). In contrast, complementation with the mutants restored the percentage of chromatid exchanges to control and WT-BRCA1 levels (Fig 6G & I).

Taken together the S114-P115 region contributes support to nascent DNA protection, the prevention of chromosome breakage and HU resistance. The data are consistent with a model in which *trans*-BRCA1 promotes replication fork protection thereby preventing DNA breaks and cell death following replication stress. Radials are also associated with short-term HU treatment in cells lacking BRCA2 or RAD51-paralogs ^8,62^. They are a result of illegitimate recombination, and may represent toxic-NHEJ of cells deficient in HR-repair^63^. We find they correlate with proficient response to Olaparib and are not associated with the fork protection defect associated with the phosphorylation-isomerisation region of BRCA1.

### Patient variants define novel functional regions of BRCA1-BARD1 required for fork protection

Our data are surprising in having established a region of BRCA1 required for the protection of replication forks and the prevention of replication-stress mediated genome instability and cell death that has no role in PARPi sensitivity and shows no features of promoting HR. We were therefore interested to know if genetic changes identified in patients with a family or personal history of breast or ovarian cancer generate mutant BRCA1 protein with similar features. The Breast cancer information core, cBioportal and literature shows genetic variants that alter the coding sequence within or close to the BRCA1 S114-P115 phosphorylation-isomerisation site and the BARD1-RAD51 binding site (https://research.nhgri.nih.gov/bic/ ^64,65^ ^66^). We generated point mutations corresponding to those that introduce amino acid changes with high Grantham difference and low variance (i.e are chemically highly distinct from the reference residue and occur in a residue highly conserved across species) ^67^ (Fig 7A-B). *BRCA1* patient variants, S114P, R133C, Y179C, S184C, S256Y showed shorter CldU tract lengths consistent with a fork protection defect (Fig 7C-G). *BARD1* patient variants located in the RAD51 binding domain of BARD1, with the exception of D135Y, also showed shorter CldU tract lengths compared to controls (Fig 7H-I). Moreover all mutants that showed a fork protection defect, also showed increased sensitivity to HU, but not to Olaparib, in colony survival assays (Fig 7J-M and Supplementary Fig 7A-D). Intriguingly Y101N-BRCA1, which is proficient for fork protection and shows HU resistance, is sensitive to PARPi, demonstrating functional separation in the other direction (Fig 7E, G, L and Supplementary Fig 7C). These data confirm the identification of novel regions of the heterodimer, mutated in cancers, that are required for fork protection and response to replicative stress that do not promote cellular responses to PARP-inhibitor, Olaparib.

**Figure 7.**
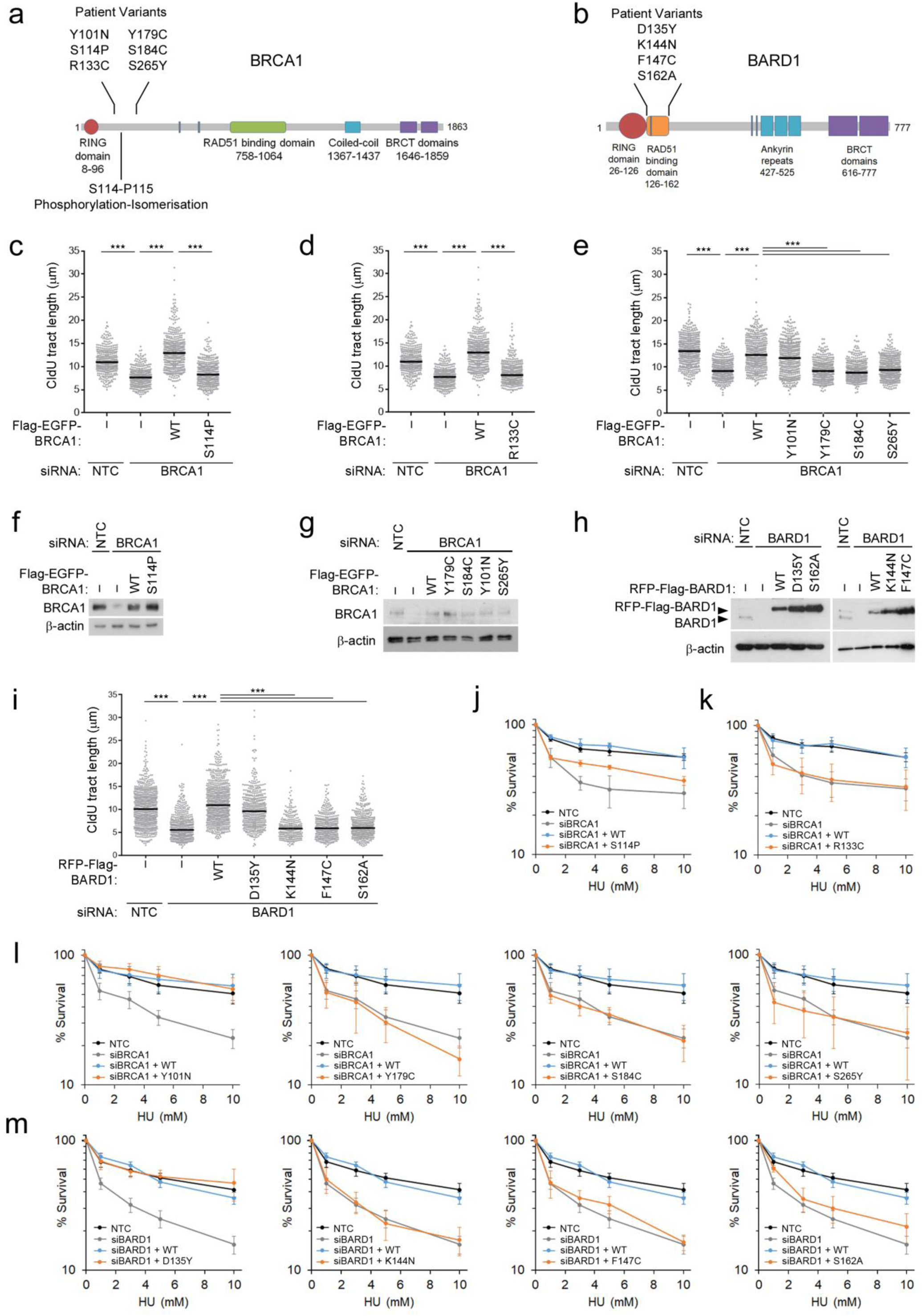
Patient variants define novel functional regions of BRCA1-BARD1 required for fork protection. a. Schematic showing approximate location of the BRCA1 S114-P115 phosphorylation-isomerisation site in full length BRCA1 and identification of patient variants proximal to this site that have a high class 65 Grantham score. b. Schematic showing approximate location of the BARD1 RAD51 binding domain and the identification of patient variants within this region that have a high class 65 Grantham score. c. CldU fibre tract lengths were measured from U20S cells depleted for BRCA1 and complemented with Flag-EGFP-BRCA1 S114P patient variant and treated with 5 mM HU for 3 hours. N>**=**340 fibres from 3 independent experiments, bars indicate median. d. CldU fibre tract lengths were measured from U20S cells depleted for BRCA1 and complemented with Flag-EGFP-BRCA1 R133C patient variant and treated with 5 mM HU for 3 hours. N>**=**340 fibres from 3 independent experiments, bars indicate median. e. CldU fibre tract lengths were measured from U20S cells depleted for BRCA1 and complemented with Flag-EGFP-BRCA1 patient variants as indicated and treated with 5 mM HU for 3 hours. N=400 fibres from 3 independent experiments, bars indicate median. f. Western blot to show BRCA1 depletions and complementation for C. g. Western blot to show BRCA1 depletions and complementation for E. h. Western blot to show BARD1 depletions and complementation for I. i. CldU fibre tract lengths were measured from U20S cells depleted for BARD1 and complemented with RFP-Flag-BARD1 patient variants as indicated and treated with 5 mM HU for 3 hours. N>300 fibres from 3 independent experiments, bars indicate median. j. Colony survival following 16 hour treatment with HU was measured in HeLa cells depleted for BRCA1 and complemented with WT or Flag-EGFP-BRCA1 S114P patient variant as indicated. N=7, error bars are SEM. k. Colony survival following 16 hour treatment with HU was measured in HeLa cells depleted for BRCA1 and complemented with WT or Flag-EGFP-BRCA1 R133C patient variant as indicated. N=3, error bars are SEM. l. Colony survival following 16 hour treatment with HU was measured in U20S cells depleted for BRCA1 and complemented with WT or Flag-EGFP-BRCA1 patient variants as indicated. N=3, error bars are SEM. m. Colony survival following 16 hour treatment with HU was measured in U20S cells depleted for BARD1 and complemented with WT or RFP-Flag-BARD1 patient variants as indicated. N=4, error bars are SEM.

**Figure 8.**
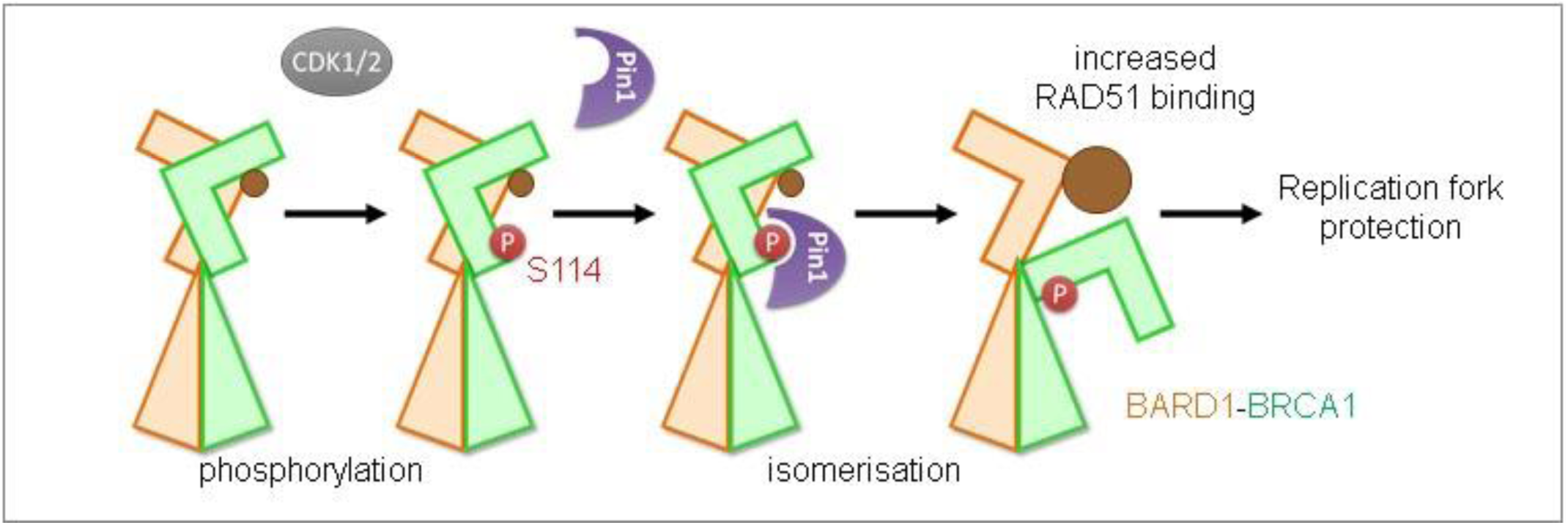
PIN1 isomerisation of BRCA1 promotes RAD51 association and replication fork protection. 1. Schematic to illustrate CDK1/2 (grey) phosphorylation at S114 (red) and subsequent PIN1 (purple) isomerisation events on the BRCA1 (orange) and BARD1 (green) N-termini. BRCA1 isomerisation (*trans*BRCA1) enhances the ability of BARD1 to associate with RAD51 (brown) and this enhanced binding is required for replication fork protection.

## Discussion

Our study reveals that CDK-PIN1 regulated conformational change in BRCA1 results in a new interface between BRCA1-BARD1 and RAD51 to promote fork protection. While PIN1 activity is capable of disrupting dimers and aggregates ^68^ ^69^ and of driving interactions specific for *cis* or *trans* conformations (reviewed in ^70^), the mechanism we identify here, that of isomerisation on one partner of a heterodimer, driving improved protein:protein interaction largely mediated by the other heterodimer partner, appears unique. We do not rule out the creation of new contacts on both BRCA1 and BARD1, indeed the cancer-associated patient mutations we identify extend the regions of both proteins involved in fork protection. We speculate these mutations represent sensitivities of a new domain structure brought about by proline isomerisation at BRCA1-P115.

In addition we reveal post-translational modifications induced by HU-treatment are required for fork protection. The CDK-PIN1 regulation of BRCA1 dove-tails with that of BRCA2 regulation in which CDK2-mediated phosphorylation of BRCA2, which would inhibit its binding to RAD51, is actively repressed in S-phase by ATR signalling and the components of the core Hippo pathway ^71,72^.

Our findings build on the view that a greater stability of RAD51-ssDNA is needed for fork protection. We show *trans* BRCA1-BARD1 specifically increases the degree of direct RAD51 binding. Since BRCA1 fork protection and fork restart processes are not coupled (Fig2A & F), whether the stability provided by BRCA1-BARD1-RAD51 interaction is relevant to this process remains to be investigated.

The separation of function mutation at BRCA1-S114A, is particular in exhibiting a fork protection defect but having no measurable impact on fork stalling, fork restart or HR and thus in examining chromosome aberrations we identify a close correlation between poor fork protection and chromosomal breaks. How breaks occur is not clear but they may arise from the processing of stalled, de-protected structures to allow cells to survive into mitosis (eg _73_). We expect that our analysis of metaphases after short-term HU under-represents chromosome damage since unprotected forks can also be converted into anaphase bridges and subsequently broken later in mitosis ^19,74^.

Our data reveals a direct function for BRCA1 in fork protection that is separate from the canonical PALB2-BRCA2 recruitment, leaving the question of how PALB2 and BRCA2 are recruited to stalled forks? In this context it may be relevant that PALB2 can be recruited through RNF168 or phosphorylated RPA at DNA double-strand breaks and both are present at stalled replication forks^75-77^.

Such an elaborate mechanism of BRCA1-BARD1 regulation, specific to fork protection, also raises the question of why RAD51 interaction is so exquisitely regulated. One possibility is that excessive RAD51 can be deleterious. Indeed excessive RAD51 at replication forks results in fork collapse in unperturbed replication^20,78,79^ and breast and lung cancer patients with high levels of RAD51 tend to do poorly compared to patients with lower levels ^12^.

We report eight patient-derived variants in BRCA1-BARD1 that impair fork protection, but not _HR_. These include a germline patient variant corresponding to the S114 phosphorylation site, *BRCA1 T459Ca,c* (pS114P) in *BRCA1* exon 7 from a 35 year old patient from northern India with stage II, lymph node positive familial breast cancer^66^. The impact of these mutants implies a wider role for fork protection in cancer development than previously described^16,17^. Whether other regions of BRCA1-BARD1 are also specifically required for fork protection, and whether loss of fork protection is sufficient to result in cancer predisposition requires further investigation.

PIN1 is amplified or highly expressed in ~15% of common cancers ^80-82^). It has not previously been implicated in endogenous mammalian DNA replication, although indirectly elevated expression is likely to increase replicative stress through oncogene activation and the induction of a shortened G1 phase ^83-89^. Its up-regulation acts to potentiate the function of several oncogenic pathways driving cell cycle progression and cell proliferation ^43,85,90-93^. Several PIN1 small molecule antagonists and peptide inhibitors have been described (reviewed in ^94^). Recently all-trans retinoic acid, used in the treatment of acute promyelocytic leukemia, was found to result in PIN1 degradation and thus the application of the drug a to wider cancer repertoire suggested ^95,96^. Our finding that PIN1 is required for replication fork protection reveals a new cross-talk pathway that may bring opportunities in cancer therapies, potentially improving the efficacy of replication stressing or immune-checkpoint therapeutics by increasing cytotoxicity and mutagenic load.

## Materials and Methods

### Tissue culture

Flp-In™ Hela, U20S and HEK293 cells were grown in Dulbeccos Modified Eagle Media (DMEM) supplemented with 10% Fetal Bovine Serum (FBS) and 1% Penicillin/Streptomycin. Cells were cultured in Corning T75 flasks and 10 cm^2^ plates and kept at 5% CO_2_ and 37°C. Once cells reached 70-80% confluency they were passaged. Cells were tested for Mycoplasma by Hoescht staining.

### Inducible stable cell line generation

Flp-In™ HeLa, U20S and HEK293 cells were plated in 10 cm^2^ dishes and transfected with a mixture containing the gene of interest cDNA in the pcDNA5/FRT/TO vector and the Flp recombinase cDNA in the pOG44 vector. Control transfections were carried out without the pOG44 recombinase. Two days after transfection, cells were preselected with 100 μg/ml Hygromycin, cell culture medium was replaced every 2-3 days and cells were selected for approximately 2 weeks. After selection cells were expanded and tested for expression of Flag-EGFP-BRCA1, RFP-Flag-BARD1 or Flag-PALB2. Cells were treated with 2 μg/ml Doxycycline for 24, 48 and 72 hours and expression levels were checked by western blotting.

### Plasmid and siRNA transfection

FuGENE 6 (*Roche*) was used as a reagent to transfect DNA plasmids into cells, the ratio used was 3:1 FuGENE (μl): DNA (μg), following the manufacturer’s guidelines. siRNA transfections were carried out using the transfection reagent Dharmafect1 (*Dharmacon*) following the manufacturer’s instructions. For a full list of siRNA sequences see Supplementary Table 1.

### Colony survival assays

Flp-In™ U20S or Hela cells were plated in 24 well plates at 4 × 10^4^ cells/ml and treated according to the experiment performed. Cells were trypsinised in 100 μl of 1x Trypsin and resuspended in 900 μl of PBS. Cells were plated out at limiting dilutions and incubated for a further 10-14 days at 37 °C at 5 % CO_2_. Once colonies had grown they were stained with 0.5 % Crystal violet in 50 % methanol and counted.

### Metaphase spreads

Flp-In™ U20S cells were treated with 5 mM HU for 4 hours and then incubated with Colcemid (0.05 µg/ml) 16 hours. Cells were trypsinized and centrifuged at 1200 rpm for 5 minutes. Supernatant was discarded and cells resuspended in PBS. 5 ml of ice-cold 0.56 % KCl solution was added and incubated at room temperature for 15 min before centrifuging at 1200 rpm for 5 min. Supernatant was discarded and cell pellet broken before fixation. Cells were then fixed in 5 ml of ice-cold methanol: glacial acetic acid (3:1). Fixation agents were removed and 10 μl of cell suspension was dropped onto alcohol cleaned slide. Slides were allowed to dry at least 24 hours and then stained with Giemsa solution (Sigma) diluted 1:20 for 20 min. Slide mounting was performed with Eukitt (Sigma).

### Proximity Ligation Assay

Flp-In™ U20S cells were seeded at a 4 × 10^4^ cells/ ml confluence onto poly-L-lysine coated coverslips and EdU pulsed for 10 minutes at 37°C for 10 minutes. 5 mM HU was then added into media for 4 hours at 37 °C. Cells were pre-extracted for 5 minutes on ice with Pre-extraction buffer (20 mM NaCl, 3 mM MgCl_2_, 300 mM Sucrose, 10 mM PIPES, 0.5 % Triton X-100) and fixed in 4 % PFA for 10 minutes before blocking in 3 % BSA for 16 hours. Blocking media was removed and click it reaction cocktail (PBS, 10 μM Biotin Azide, 10 mM sodium ascorbate, 1 mM CuSO_4_) was added for 1 hour at room temperature. Click it reaction was washed and cells blocked in 5 % BSA for 30 minutes. Cells were then incubated with primary antibodies, Biotin (Jackson Immunoresearch) and RAD51 (Calbiochem) in 3 % FCS in PBS for 1 hour at room temperature. After incubation with primary antibodies cells were incubated with the MINUS/PLUS PLA probes (Sigma Duolink PLA kit) for 1 hour at 37 °C in a warm foil covered box. Cells were then washed twice for 5 minutes with wash buffer A (Sigma Duolink PLA kit) and incubated with the Sigma Duolink Ligation kit (1X ligation buffer, ligase enzyme) for 30 minutes at 37 °C. Cells were washed twice for 5 minutes with wash buffer A and incubated for 100 minutes at 37 °C with the Sigma Duolink amplification kit (1X amplification buffer, polymerase enzyme). Finally they were washed twice for 10 minutes with wash buffer B at room temperature and coverslips were mounted using the Duolink mounting media with DAPI (Sigma).

### GST-PIN1 Pull down assay

Cells were washed with 10 ml ice cold PBS before being lysed in 5 ml TG lysis buffer (40 mM Tris-HCL pH 8, 274 mM NaCl, 2 mM EGTA, 3 mM MgCl_2_, 2 % Triton x100, 20 % Glycerol) with addition of cOmplete protease inhibitor cocktail (Roche) and PhosSTOP (Roche) tablets. The lysed cells were transferred into 1.5 ml Eppendorf tubes incubated on ice for 20 mins and sonicated twice at 20 % intensity for 10 seconds. Samples were spun at 13000 rpm at 4 °C for 10 mins and the supernatant kept. 50 μl of the supernatant was mixed with 20 μl 4x SDS Loading buffer and boiled at 95 °C for 5 mins. 800 μl of the cell supernatant was then incubated with equal concentrations of GST-WW PIN1 and GST-W34A PIN1 beads for 2 hours at 4°C. The GST-pull downs were washed three times in TG lysis buffer before adding 60 μl 4x SDS loading buffer directly to the beads. Samples were boiled at 95°C for 5 mins and then 40 μl loaded onto an SDS PAGE gel and analyzed by western blotting.

### Flag immunoprecipitation

Cells were plated in a 10 cm^2^ plate and treated with Doxycycline for 48 hours to express inducible EGFP-Flag-BRCA1 or RFP-Flag-BARD1. Cells were washed with 10 ml ice cold PBS before being scraped in ice cold Nuclear Lysis Buffer (10 mM HEPES pH7.6, 200 mM NaCl, 1.5 mM MgCl_2_, 10 % Glycerol, 0.2 mM EDTA, 1 % Triton) with addition of cOmplete protease inhibitor cocktail (Roche) and PhosSTOP (Roche) tablets and 20 μM MG132 for every 10 ml. The lysed cells were then transferred into 1.5 ml Eppendorf tubes incubated on ice for 10 mins, and sonicated 1 time at 20% intensity for 10 seconds. Samples were spun at 13000 rpm at 4°C for 10 mins and the supernatant kept, the pellet was discarded. 50 μl of the supernatant was mixed with 20 μl 4x SDS Loading buffer and boiled at 95°C for 5 mins. For every IP, 10 μl Flag-agarose beads were firstly washed out of storage buffer by doing 3x 1ml PBS washes and centrifuging at 3000 rpm between each wash. 90 μl of PBS was added for every 10 μl of agarose beads. Once the beads were resuspended in PBS, 100 μl were transferred into an Eppendorf with 500 μl of supernatant and 500 μl of PBS. The Eppendorfs were rotated for 2 hours at 4°C. Samples were centrifuged at 3000 rpm for 1 min and the beads left to settle. The supernatant was then removed before 3x 1 ml PBS-0.02% tween washes. The wash buffer was completely removed before adding 60 μl 2x SDS loading buffer. This was boiled at 95°C for 5 mins and then 20 μl loaded onto an SDS PAGE gel and analysed by western blotting.

### Fibre Labelling and spreading

Cells were seeded in 6 cm^2^ plates at a density of 20x 10^4^ cells/well and treated with thymidine analogues. To monitor stability of nascent DNA cells were incubated at 37 °C with CldU for 20 mins at a final concentration of 25 μM and then with 5 mM HU for 3 hours. To monitor replication fork restart cells were incubated at 37 °C with CldU for 20 mins at a final concentration of 25 μM and then with 5 mM HU for 3 hours. The HU was then washed out with 3x PBS washes and cells were incubated for a further 40 mins in media containing 250 μM IdU at 37°C. After incubation with thymidine analogues, cells were washed 2x with ice-cold PBS for 5 minutes with rotation then trypsinised, resuspended in 1ml of PBS and counted. The optimal concentration is 50 × 10^4^ cells/ml and thus cells were adjusted to such concentration. 2 μl of the cell sample was placed on Snowcoat microscope slides and allowed to slightly dry for 7 mins. Then 7 μl of spreading buffer (200 mM Tris pH7.4, 50 mM EDTA, 0.5% SDS) was mixed with the sample and incubated for 2 mins to lyse the cells. In order to spread the sample down the slide, slides were gradually tilted and once the sample had reached the bottom of the slide, they were allowed to dry for 2 mins. Finally, slides were fixed in a 3:1 ratio of Methanol: Acetic acid for 10 mins before leaving slides to air dry for 5-10 mins. Dried slides were stored at 4°C till staining.

### Fibre Immunostaining

After fibre spreading slides were washed twice for 5 minutes with 1 ml H_2_O and rinsed with 2.5 M HCl before denaturing DNA with 2.5 M HCl for 1 hour 15 mins. Slides were then rinsed 2 × with PBS and washed for 5 minutes in blocking solution (PBS, 1 % BSA, 0.1 % Tween20). Slides were incubated for 1 hour in blocking solution. After blocking, each slide was incubated with 115 μl of primary antibodies, Rat αBrdU (AbD Serotec/Abcam) used at a concentration of 1:2000 and Mouse αBrdU (Becton Dickinson) used at 1:750. Slides were covered with large coverslips and incubated with the antibodies for 1 hour. After incubation with the primary antibody, slides were rinsed 3x with PBS and then incubated for 1 min, 5 mins and 30 mins, with blocking solution. After rinsing and washing, slides were incubated with 115μl of secondary antibodies (α-Rat AlexaFluor 555 and α-Mouse AlexaFluor 488) in blocking solution, at a concentration of 1:500, covered with a large coverslip for 2 hours. Slides were rinsed 3x with PBS and incubated with blocking solution for 1 min, 5 mins and 30 mins. After again rinsing 2 × with PBS, immunomount mounting media was added to the slide and a large coverslip placed over and left to dry. Coverslips were then stored at −20°C for microscopy analysis. It is important to point out that during this process slides must be kept protected from light.

### Immunofluorescent staining

U20S WT or S114A-BRCA1 cells were plated at a density of 5 × 10^4^ cells/ml in 24 well plates on circular glass coverslips (13 mm). Cells were treated siBRCA1 for 48 hours and complemented with WT or S114A BRCA1 by addition of Doxycycline. Cells were then treated with 20 µM Olaparib for 2 hours. Cells were pre-permeablised by incubation with 0.5 % Triton-PBS on ice for 10 minutes before fixation with 4 % PFA. Once fixed the cells were permeabilised for a further 5 mins using 0.5 % TritonX in PBS before incubation with blocking solution - 10% FCS in PBS for 30 mins. Cells were then incubated with rabbit polyclonal RAD51 (H92) antibody in 10% FCS PBS at room temperature overnight. The following day, cells were washed in PBS/FCS before incubating them with AlexaFluor- 488 secondary antibody at a 1:2000 concentration, for 2 hours. Cells were then washed three times in PBS and the DNA stained using Hoescht at a 1:20000 concentration for 5 mins. Excess of Hoescht was washed with PBS and coverslips mounted onto Snowcoat slides using Immunomount mounting media.

### EdU staining

Cells subjected to immunofluorescent staining were also subjected to EdU staining. Cells were incubated with 10 μM final concentration of EdU for 2 hours prior to fixing and staining was carried out as detailed in the Click-iT® EdU Imaging Kits (Life Technologies).

### CldU Immunostaining

Cells were plated at a density of 5 × 10^4^ cells/ml in 24 well plates on circular glass coverslips (13 mm). Cells were treated as described and incubated at 37 °C with CldU for 20 mins at a final concentration of 25 μM. Cells were then fixed with 4 % PFA and permeabilised for 5 mins using 0.25 % TritonX in PBS. After permeabilisation, cells were washed twice with PBS and twice with blocking solution containing 10 % FCS in PBS. 2 M HCl was added for 30 mins at 37 °C and cells incubated with 10 % FCS in PBS for 1 hour at room temperature (RT). Cells were then incubated with FLAG (M2-Sigma)(1:1000) and Rat αBrdU (AbD Serotec) (1:500), in 10 % FCS PBS at RT, for 1 hour. After primary antibody incubation, cells were washed 3x in PBS and then incubated for 10 mins in stringency buffer (0.5 M NaCl, 36 mM Tris pH 7.5-8, 0.5 % Tween20), before incubating them with AlexaFluor antbodies at a 1:500 concentration, for 2 hours. Cells were then washed twice with PBS and once with stringency buffer, and DNA stained using Hoescht at a 1:20000 concentration for 5 mins. Excess of Hoescht was washed with PBS and coverslips mounted onto Snowcoat slides using Immunomount mounting media.

### Microscopy

Immunofluorescent staining was imaged using the Leica DM6000B microscope using a HBO lamp with 100W mercury short arc UV bulb light source and four filter cubes, A4, L5, N3 and Y5, which produce excitations at wavelengths 360 488, 555 and 647 nm respectively.

### Western blotting

Acrylamide SDS PAGE protein gels were run and then transferred onto PVDF Immobilon-P membrane, which was previously activated with methanol. Transfers were set up in Biorad Trans-blot cassettes with a sponge and 2 pieces of filter paper either side of the membrane and gel. The cassette was then placed in the tank with 1x Transfer buffer 20 % Methanol and ran at 100 volts for 1 hour.

After the transfer the membrane was blocked in 5 % marvel milk in PBS with 0.1 % Tween (PBStw) or in 5 % BSA with PBStw, for a minimum of 1 hour. Membranes were then incubated in primary antibody (for a full list of antibodies see Supplementary Table 2) accordingly at 4 °C on a roller. Blots were then washed 3x 10 mins in PBStw and then transferred into secondary HRP antibodies in 5 % marvel milk for a minimum of 1 hour whilst being rocked. After secondary incubation, blots were again washed 3x 10 mins in PBStw. Finally membranes were probed with 1:1 EZ-ECL mix (Biological Industries), the excess of ECL was removed and the membrane was placed into a plastic wallet that was placed inside a cassette. A Fuji film X-Ray film was placed in the cassette for differential length of time and then it was developed using the Xograph Compact X4 developer. Densitometry calculations were performed using Image J.

### p-S114 Antibody Generation

Custom mouse monoclonal and rabbit polyclonal antibodies were raised against BRCA1 phospho-S114 by Genscript using the following peptide: CFAKKENNpSPEHLKD

### BRCA1:BARD1 protein expression and purification

#### WT and P115A His-BRCA1_1-500_-BARD127-327 for in vitro analysis

The expression of His-BRCA1-WT + BARD1-WT and His-BRCA1-PA + BARD1-WT in Rosetta™(DE3) was induced by the addition of 1 mM Isopropyl-β-d-thiogalactopyranoside (IPTG), and the proteins were produced in LB medium containing 50 μg/ml of kanamycin, 100 μg/ml of ampicillin and 30 μg/ml of chloramphenicol at 37°C for 5 hours. For purification of the His-BRCA1-WT + BARD1-WT and His-BRCA1-PA + BARD1-WT products, the cells were harvested and resuspended in 20 mM HEPES potassium salt, pH 7.4, 50 mM Imidazole, 500 mM NaCl, 1.0 mM TCEP [tris(2-carboxyethyl)phosphine], cOmplete EDTA-free protease inhibitor cocktail tablet (Roche). Cells were lysed using an Emulsiflex-C3 homogenizer (Avestin) and broken by three passages through the chilled cell. The lysate was centrifuged at 75,000 xg using a JA 25.50 rotor (Beckman Coulter) and filtered through a 0.45-μm filter. The clarified lysate was applied onto a 5-ml HisTrap HP column (GE Healthcare). The column was washed extensively using the same buffer, and the protein was eluted using buffer containing 500 mM imidazole.

Fractions containing a band of the correct size were concentrated using a Vivaspin 20-ml concentrator (10,000 molecular weight cut-off [MWCO]) (GE Healthcare) and gel purified using an Akta Pure 25 (GE Healthcare LS) with a prepacked Hi-Load 10/300 Superdex 200 PG column.

#### WT and S114A BRCA1_1-300_ for CDK2 kinase assay

BRCA1 and BARD1 proteins were expressed from pET15b-His-BRCA1_1-300_:His-BARD1_26-142_ vector in BL21(DE3) bacteria (Bioline). Bacteria were grown at 37 °C until an optical density of 0.6 was reached. Protein expression was induced by addition of 0.5 mM isopropyl β-D-1-thiogalactopyranoside (IPTG) (Bioline), and the temperature was immediately decreased to 25 °C. Bacteria were grown for a further 24 h. Bacterial pellets were collected after centrifugation at 3,000*g* for 10 min at 4 °C and then lysed in ice-cold lysis buffer (50 mM sodium phosphate, pH 7, 300 mM sodium chloride, 5% glycerol and 10 mM β-mercaptoethanol). Lysates were sonicated for 1 min at 30 % intensity and then clarified by centrifugation at 14000*g* for 10 min at 4 °C. The supernatant was incubated with 0.25 ml His-select beads (Sigma) overnight at 4 °C with rotation. The following day, the beads were washed three times with ice-cold wash buffer (50 mM sodium phosphate, pH 7, 300 mM sodium chloride, 5 % glycerol, 10 mM β-mercaptoethanol and 50 mM imidazole) before elution on ice in 50 mM sodium phosphate, pH 7, 300 mM sodium chloride, 5 % glycerol, 10 mM β-mercaptoethanol and 300 mM imidazole. Purified proteins were dialyzed against (25 mM Tris-HCl, pH 7.5, 10 % glycerol, 2 mM dithiothreitol (DTT) and 150 mM potassium chloride), and purity was assessed by resolution on a 15 % SDS–PAGE gel.

### RAD51 in vitro binding assay

0.5 μl of Human recombinant RAD51 (Abcam) was incubated with 40 μl of a 50% slurry of His-BRCA1-WT + BARD1-WT or His-BRCA1-PA + BARD1-WT immobilized in Ni^2+^-resin together with 500 μl of RAD51 binding buffer (25 mM Tris-HCl pH 7.5, 10% Glycerol, 0.5 mM EDTA, 0.05% Igepal CA-630, 1mM 2-mercaptoethanol, 150 mM KCl, 50mM Imidazole) for 30 minutes at 4°C in rotation. After incubation the resin was washed three times with the RAD51 binding buffer before eluting in 40 μl 4x SDS loading buffer at 95°C for 5 minutes. The SDS elute was then analyzed by Western blotting and Coomassie blue staining.

### Trypsin proteolytic digestion

The His-BRCA1_1-500_-BARD1_27-327_ fragment was subjected to trypsin proteolytic digestion and 50 μl of the His-BRCA1-WT + BARD1-WT or the His-BRCA1-PA + BARD1-WT (0.5 mg/ml) was incubated with 50 μl of the Trypsin digestion buffer (25 mM Tris pH 7.5, 10 % Glycerol, 2 mM DTT and 150 mM KCl) and 2 μl of trypsin (20 ng/μl). Once trypsin was added, samples were immediately incubated at 37 °C and time points collected after 3, 7, 15, 30, 60 and 90 minutes and eluted in 4x SDS-PAGE loading buffer. Samples were boiled at 95°C for 5 mins and then 5 μl loaded onto an SDS PAGE gel and analyzed by western blotting using the MS110 BRCA1 antibody.

### CDK2-Cyclin A Kinase assay

50 ng of WT or S114A His-BRCA1_1-300_:His-BARD1_26-142_ were incubated with 50ng of recombinant CDK2-Cyclin A (Sigma C0495) in 25 mM MOPS pH7.2, 12.5 mM β-glycerolphosphosphate, 25 mM magnesium chloride, 5mM EGTA, 2 mM EDTA, 0.5 mM DTT and 50 ng/µl BSA. The kinase reaction was started by addition of 10 mM ATP and samples were incubated at 30 °C for 30 minutes. The reaction was stopped by addition of 4x SDS-PAGE loading buffer and incubated at 95 °C for 5 minutes. 5 µl of the total reaction was run on 14 % SDS PAGE gel before transfer to Immobilon-P membrane (MERCK-Millipore) and western blotting for phospho-S114 (3C10G8) or BRCA1 (MS110).

### Recombinant PIN1 protein expression

Bl21 Escherichia Coli were transformed with the pGEX protein expression vector containing the protein of interest. Colonies were picked and grown up in 50 ml starter cultures containing Ampicillin at 37°C for 16 hours at 200 rpm. Starter cultures were transferred to 500 ml Luria Bertani (LB) containing Amp and grown for 2 hours at 37 °C at 200 rpm. Bacterial expression was induced using 0.5 mM IPTG and bacteria left to grow for 5 hours at 37°C at 200 rpm.

### Recombinant PIN1 Protein purification

Bacteria were pelleted by centrifuging at 12000 rpm for 10 mins at 4°C. Then the bacterial pellet was lysed in 10 ml GST lysis buffer (20 mM Tris-HCL pH8, 130 mM NaCl, 1 mM EGTA, 1.5 mM MgCl_2_, 1 % Tritonx-100, 10 % Glycerol, 1 mM DTT) with 1 protease inhibitor tablet (Roche). Bacteria were resuspended and left on ice for 20 mins and then sonicated 2 times at 20% intensity for 10 seconds. Lysed bacteria were spun 20 mins at 13000 rpm to pellet debris. Supernatant was transferred to a 50 ml falcon tube and made up to 35 ml with lysis buffer and 500 μl of pre-washed Glutathione Sepharose 4B beads (GE Life Sciences) and rotated at 4°C for 16 hours.

### Site-Directed Mutagenesis

Specific primers were designed for mutagenesis (Supplementary Table 3) and mutagenesis performed by PCR using PfU (Promega). All mutagenesis was confirmed by Sanger Sequencing (Source Bioscience).

### Statistics

All statistics were done using two-sided Student’s T-test. Significance is defined as * p<0.05, **p<0.01 and ***p<0.005 throughout.

## Author contributions

M. D-M generated constructs and cell lines, performed colony survival experiments, proximity ligation assays, fibre experiments and performed the in vitro analysis. R.M.D generated constructs and cell lines, fibre experiments and colony survival analysis. M.J generated and purified proteins and performed in vitro analysis. K.S undertook and analysed metaphase spreads. A.C performed GST-PIN1-WW analysis on BARD1 cell lines. J.B provided technical support. J.C made M1411T cell lines and performed Cisplatin survival analysis. G.S. made preliminary observations, J.R.M, R.M.D and M. D-M wrote the paper. All authors commented on the paper and ongoing research. R.M.D and J.R.M directed the project.

## Acknowledgements

Grant funding for this project was as follows. CRUK: C8820/A19062 (R.M.D. and J.B.), C17183/A23303 (GSS), Breast Cancer Now: 2015MayPR499 (K.S.). Wellcome Trust 206343/Z/17/Z (M.J.). CRUK Centre training (M. D-M.). We thank Professors Simon Cook (Babraham Inst) for PIN1 template DNA and Titia Sixma (NKI, Amsterdam) for the BARD1 construct. We also thank Dr. Philip Byrd (University of Birmingham) for useful advice regarding the PALB2 reagents, Dr. Tim Wallach for generation of the S114P mutation and Dr. Shabana Begum (University of Birmingham) for support with the proximity ligation assays. In addition, we thank the Microscopy and Imaging Services at Birmingham University in the Tech Hub facility for microscope support and maintenance.

## Data Availability

The datasets generated during the current study are available from the corresponding authors on reasonable request.

---

**Supplementary Figure 1.**
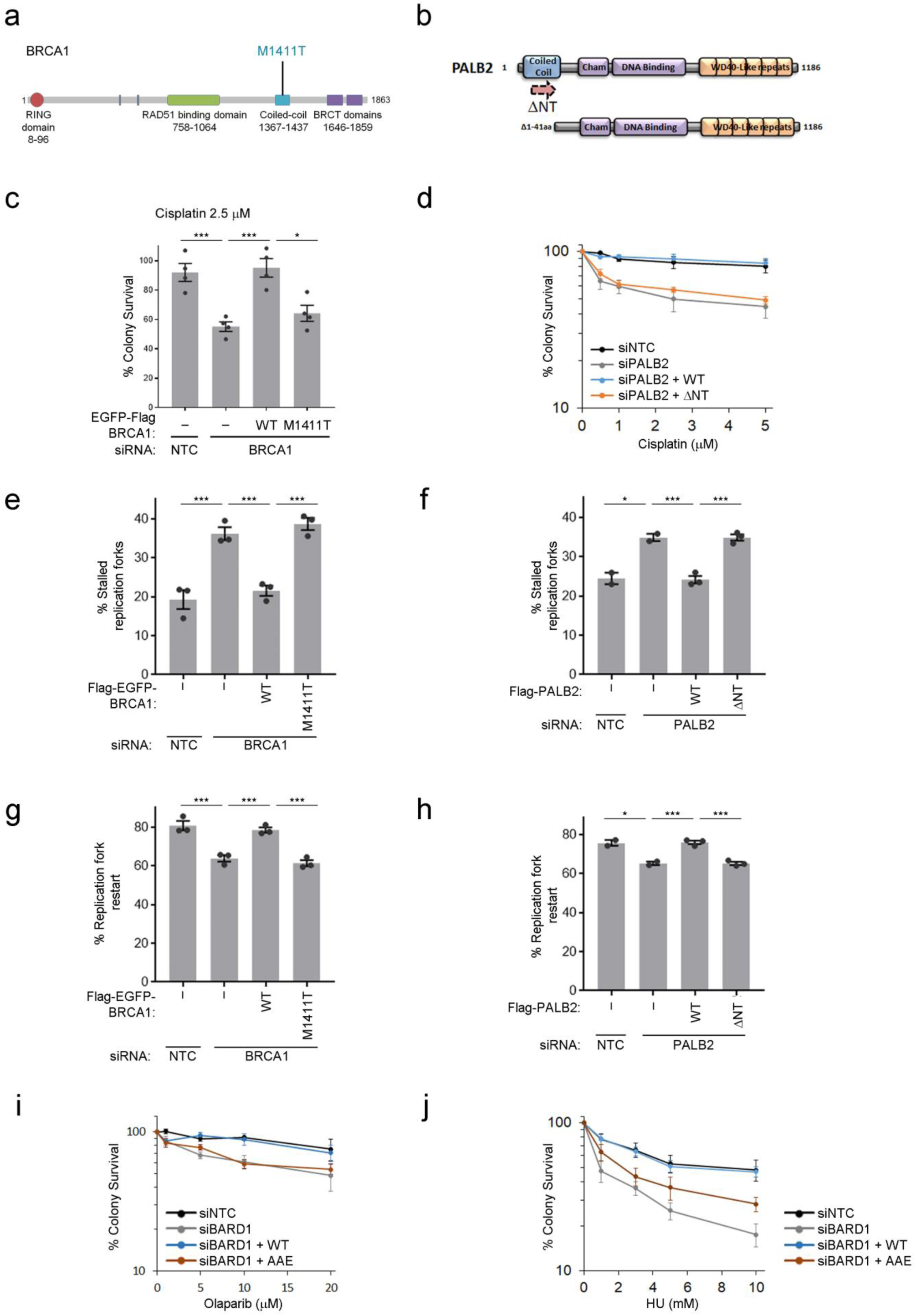
The BRCA1-BARD1 heterodimer promotes fork protection through the RAD51 binding region and not the PALB2-BRCA2 interaction region. a. Schematic of BRCA1 protein structure indicating RING (red), RAD51-binding (green), coiled-coil (blue) and BRCT repeat (purple) domains. The M1411T patient variant disrupts PALB2 binding and is located in the Coiled-coil domain. b. Schematic representation of the PALB2 protein structure indicating BRCA1-interacting coiled-coil (blue), ChAM and DNA binding (purple) and WD40-like repeat (orange) domains. The ΔNT-PALB2 mutant lacks the N-terminal Coiled-coil domain. c. Colony survival following 2 hour treatment with 2.5 µM Cisplatin was measured in HeLa cells depleted for BRCA1 and complemented with WT or M1411T Flag-EGFP-BRCA1. N=4, error bars are SEM. d. Colony survival following 2 hour treatment with Cisplatin was measured in U20S cells depleted for PALB2 and complemented with WT or ΔNT-FLAG-PALB2. N=4, error bars are SEM. e. The % stalled replication forks were measured from 3-independent experiments in U20S cells depleted for BRCA1 and complemented with WT or M1411T-Flag-EGFP-BRCA1. Grey bars indicate mean, error bars are SEM. f. The % stalled replication forks were measured from 3-independent experiments in U20S cells depleted for PALB2 and complemented with WT or ΔNT-FLAG-PALB2. Grey bars indicate mean, error bars are SEM. g. The % replication forks able to restart after release from 3 hr of 5 mM HU were measured from 3-independent experiments in U20S cells depleted for BRCA1 and complemented with WT or M1411T-Flag-EGFP-BRCA1. N=3. Grey bars indicate mean, error bars are SEM. h. The % replication forks able to restart after release from 3 hr of 5 mM HU were measured from 3-independent experiments in U20S cells depleted for PALB2 and complemented with WT or ΔNT-FLAG-PALB2. N=3. Grey bars indicate mean, error bars are SEM. i. Colony survival following 2 hour treatment with Olaparib was measured in U20S cells depleted for BARD1 and complemented with WT or AAE-RFP-Flag-BARD1. N=4, error bars are SEM. j. Colony survival following 2 hour treatment with HU was measured in U20S cells depleted for BARD1 and complemented with WT or AAE-RFP-Flag-BARD1. N=7, error bars are SEM.

**Supplementary Figure 2.**
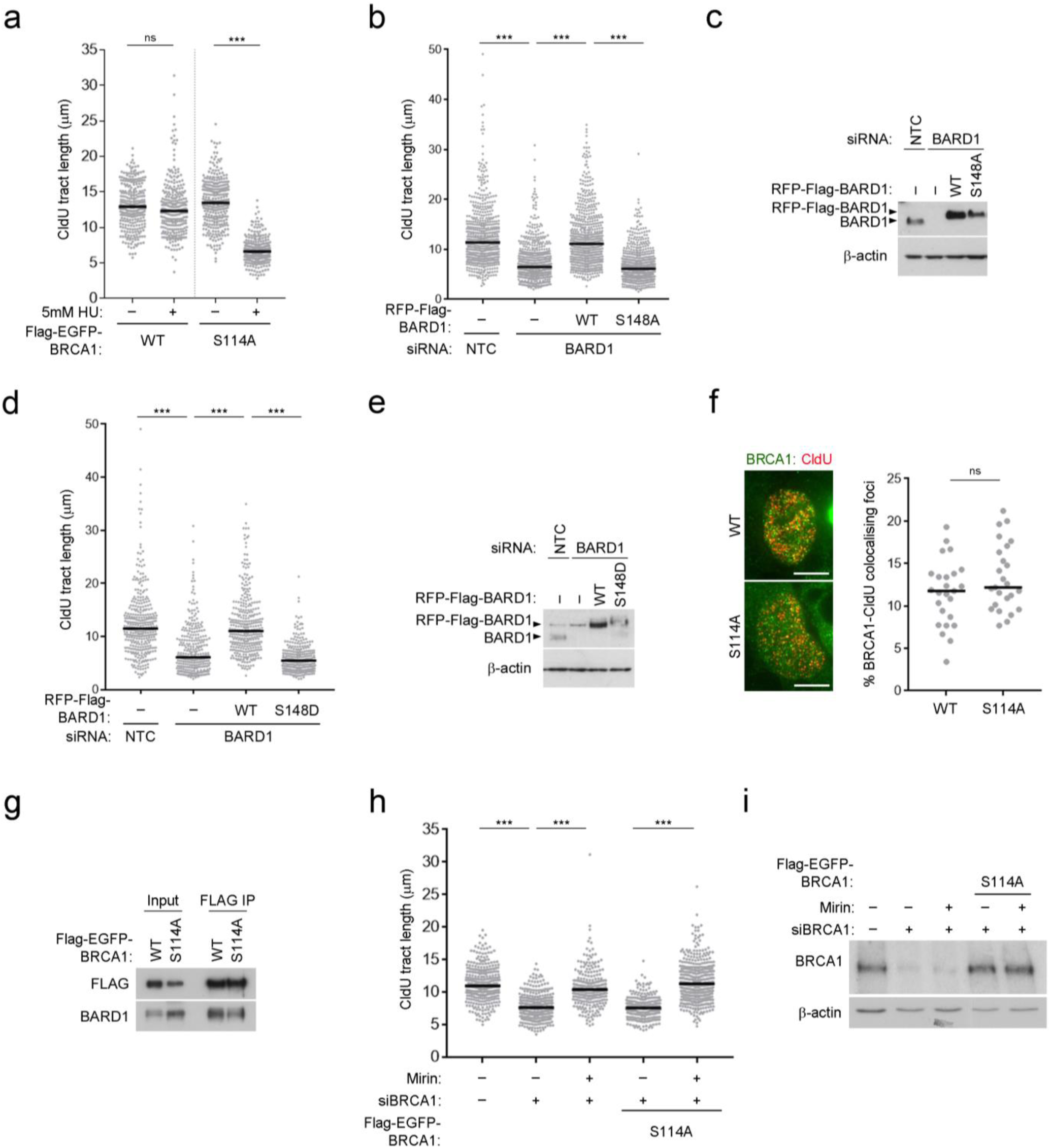
The BRCA1 serine 114 phosphorylation site is required for the protection of nascent DNA. A. CldU fibre tract lengths were measured from U20S cells depleted for BRCA1 and complemented with Flag-EGFP-BRCA1 WT or S114A and treated with or without 5 mM HU for 3 hours. N=280 fibres from 3 independent experiments, bars indicate median. B. CldU fibre tract lengths were measured from U20S cells depleted for BARD1 and complemented with RFP-Flag-BARD1 WT or S148A and treated with 5 mM HU for 3 hours. N=600 fibres from 3 independent experiments, bars indicate median. C. Western blot to show BARD1 depletions and complementation for B. D. CldU fibre tract lengths were measured from U20S cells depleted for BARD1 and complemented with RFP-Flag-BARD1 WT or S148D and treated with 5 mM HU for 3 hours. N=290 fibres from 2 independent experiments, bars indicate median. E. Western blot to show BARD1 depletions and complementation for D. F. U20S cells expressing WT or S114A Flag-EGFP-BRCA1 were pulsed with CldU before fixation and staining. The percentage of co-localising BRCA1-CldU foci per cell was scored. N=25 cells and bars indicate median. ns=not significant. G. FLAG-Immunoprecipitation of Flag-EGFP-BRCA1 from HEK293 cells showing co-purification of BARD1. H. CldU fibre tract lengths were measured from U20S cells depleted for BRCA1 and complemented with S114A-Flag-EGFP-BRCA1 and treated with or without 50 μM Mirin and HU for 3 hours. N>240 fibres from 3 independent experiments, bars indicate median. I. Western blot to show BRCA1 depletions and complementation for H.

**Supplementary Figure 3.**
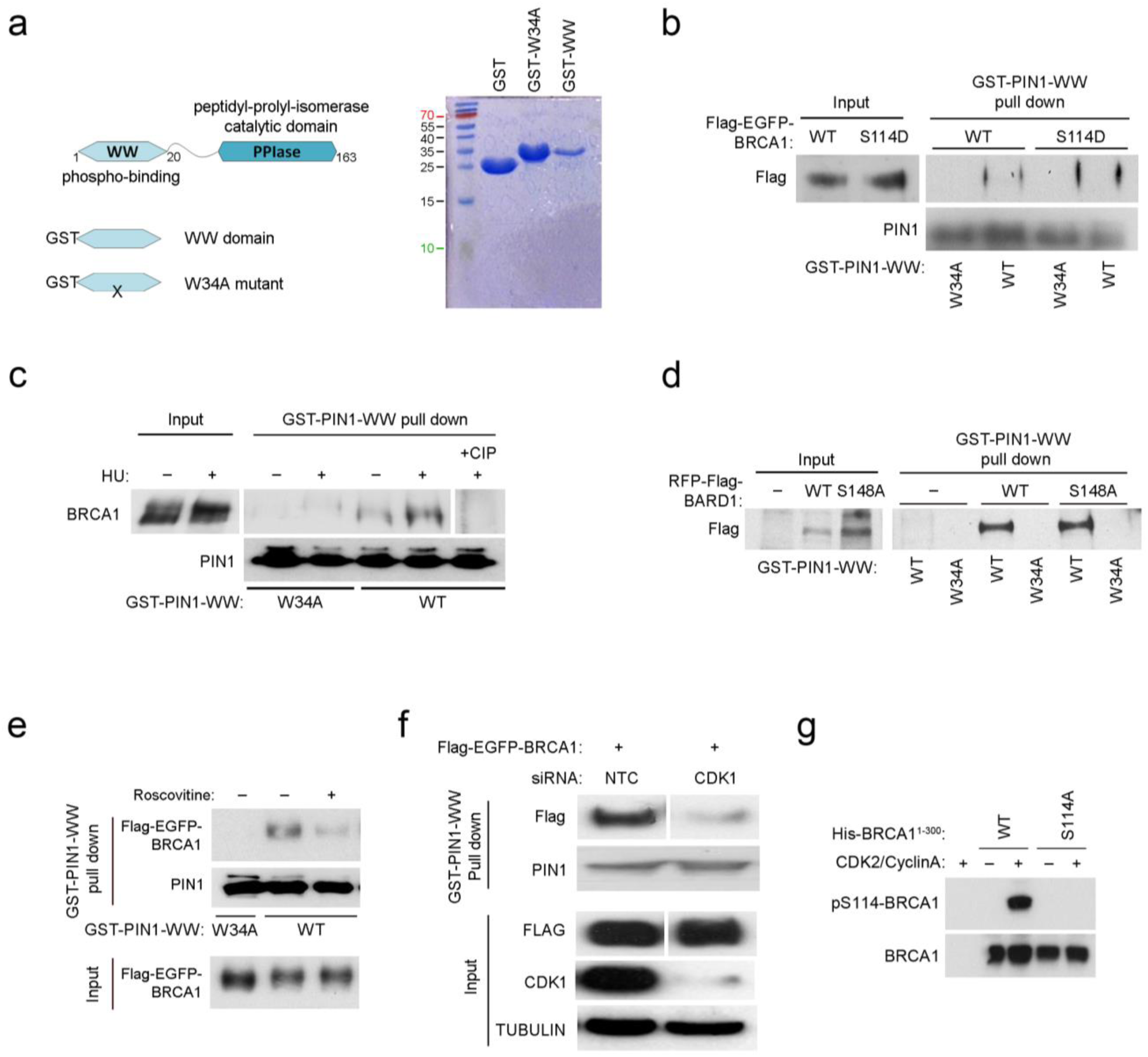
Phosphorylation of BRCA1 at serine 114 promotes PIN1 interaction. a. Schematic of the protein structure of PIN1 indicating phospho-binding WW and Peptidyl-Prolyl-Isomerase domains. Cartoons illustrate the GST-fusions of the WW domain WT and W34A phospho-binding mutant. Coomassie blots indicate recombinant GST-WW fragments purified from *E. coli*. b. Glutathione-Sepharose beads bound with the GST-fused-WW domain of PIN1 were used to pull-down WT and S114D Flag-EGFP-BRCA1, from U20S cell lysates. Beads bound by GST-W34A WW-domain were used as a negative control. c. Glutathione-Sepharose beads bound with the GST-fused-WW domain of PIN1 were used to pull-down endogenous BRCA1 from HEK293 cell lysates treated with and without 3mM HU for 6 hours. The final lane indicates lysates pre-treated with calf-intestinal phosphatase. Beads bound by GST-W34A WW-domain were used as a negative control. d. Glutathione-Sepharose beads bound with the GST-fused-WW domain of PIN1 were used to pull-down WT and S148A RFP-Flag-BARD1, from U20S cell lysates. Beads bound by GST-W34A WW-domain were used as a negative control. e. Glutathione-Sepharose beads bound with the GST-fused-WW domain of PIN1 were used to pull-down Flag-EGFP-BRCA1 from HEK293 cell lysates treated with and without 25 μM Roscovitine. Beads bound by GST-W34A WW-domain were used as a negative control. f. Glutathione-Sepharose beads bound with the GST-fused-WW domain of PIN1 were used to pull-down Flag-EGFP-BRCA1 from HEK293 cell lysates that had been depleted for CDK1. Beads bound by GST-W34A WW-domain were used as a negative control. g. Recombinant purified His-BRCA1_1-300_:His-BARD1_26-142_ were incubated with recombinant active CDK2/Cyclin A. Western blots were probed for phospho-S114-BRCA1 and BRCA1.

**Supplementary Figure 4.**
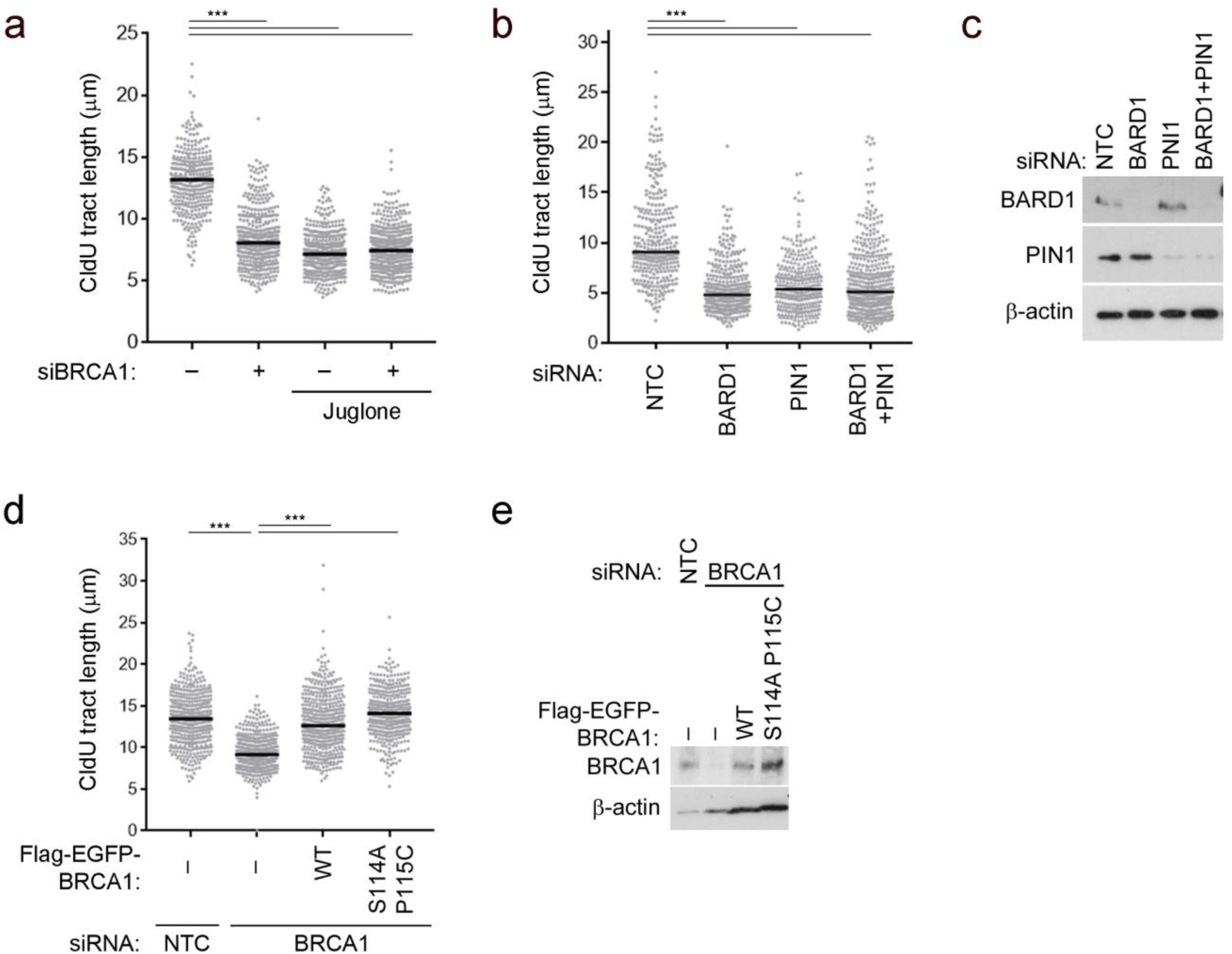
PIN1 regulates the BRCA1-BARD1 heterodimer to promote fork protection. a. CldU fibre tract lengths were measured from U20S cells depleted for BRCA1 and treated 5 mM HU for 3 hours with or without 20 μM Juglone. N=>330 fibres from 3 independent experiments, bars indicate median. b. CldU fibre tract lengths were measured from U20S cells depleted for BARD1 and/or PIN1 and treated 5 mM HU for 3 hours. N=300 fibres from 2 independent experiments, bars indicate median. c. Western blot to show BARD1 and PIN1 depletions for B. d. CldU fibre tract lengths were measured from U20S cells depleted for BRCA1 and complemented with WT or S114AP115C Flag-EGFP-BRCA1 and treated 5 mM HU for 3 hours. N=400 fibres from 3 independent experiments, bars indicate median. e. Western blot to show BRCA1 depletion and complementation as described for D.

**Supplementary Figure 5.**
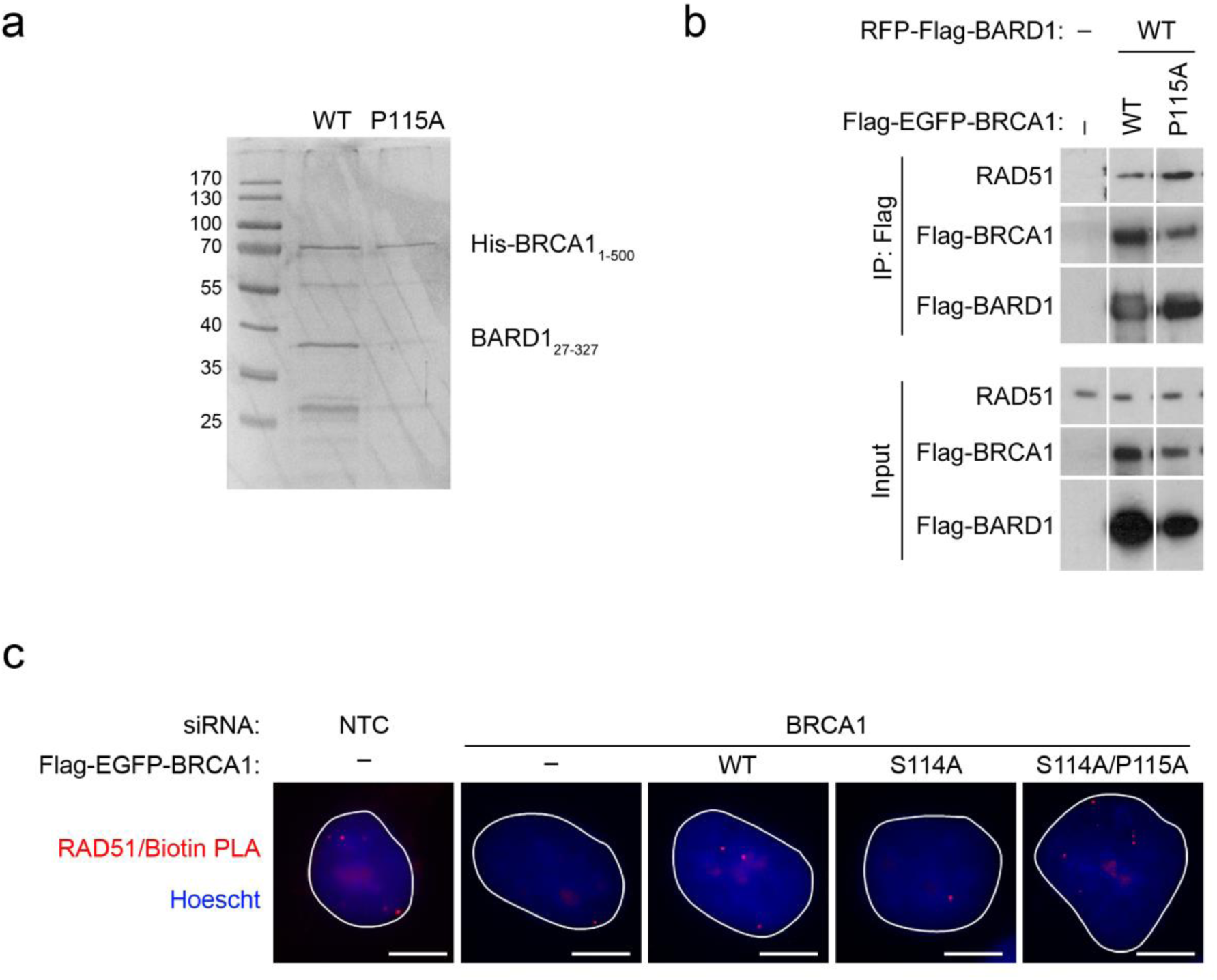
BRCA1-BARD1 isomerisation enhances direct RAD51 binding and promotes accumulation at nascent DNA. a. Coomassie gel of recombinant purified His-BRCA11-500 and BARD127-327 from *E. coli*. b. FLAG-Immunoprecipitation of Flag-EGFP-BRCA1 and RFP-Flag-BARD1 complexes from 293 cells showing co-purification of RAD51. c. Representative images for Figure 5C. RAD51 colocalisation with nascent DNA, marked by pulse labelling with EdU, was measured using the proximity ligation assay (PLA) in U20S cells depleted for BRCA1 and complemented with Flag-EGFP-BRCA1 variants as indicated. Red foci indicate RAD51/EdU-Biotin interaction in cells. Scale bars are 10 µm.

**Supplementary Figure 6.**
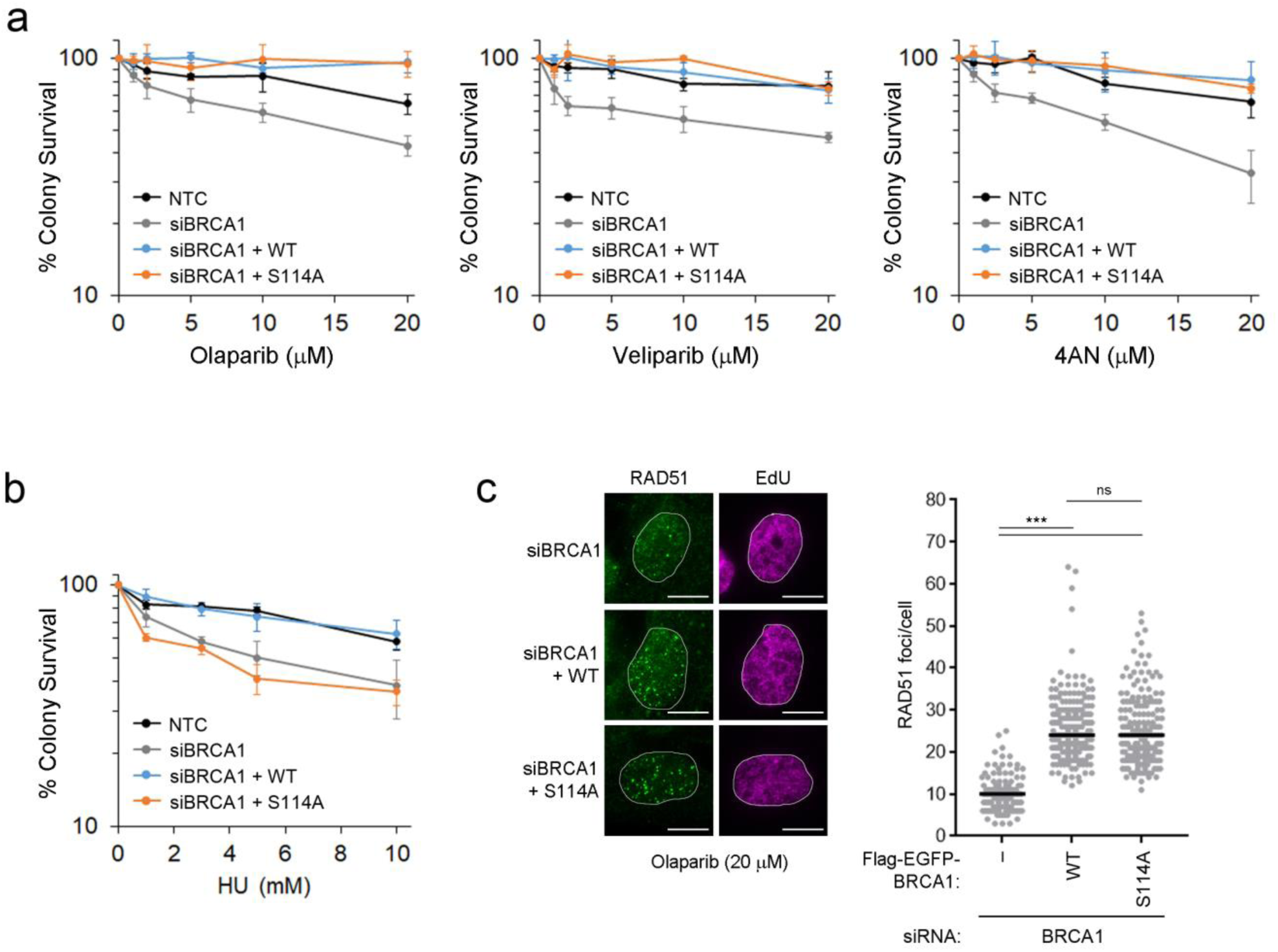
Loss of BRCA1-isomerisation leads to genomic instability and increased sensitivity to replication stress agents. a. Colony survival following 2 hour treatment with PARP inhibitors Olaparib, Veliparib and 4AN was measured in HeLa cells depleted for BRCA1 and complemented with WT or S114A Flag-EGFP-BRCA1. N>3, error bars are SEM. b. Colony survival following overnight treatment with HU was measured in U20S cells depleted for BRCA1 and complemented with WT or S114A Flag-EGFP-BRCA1. N=3, error bars are SEM. c. RAD51 foci in S-phase U20S cells marked by EdU were scored from BRCA1 depleted cells complemented with Flag-EGFP-BRCA1 WT or S114A treated with 20 µM Olaparib (2 hours). Scale bars are 10 μm. Graph shows number of RAD51 foci/EdU positive cell. Bars indicate median.

**Supplementary Figure 7.**
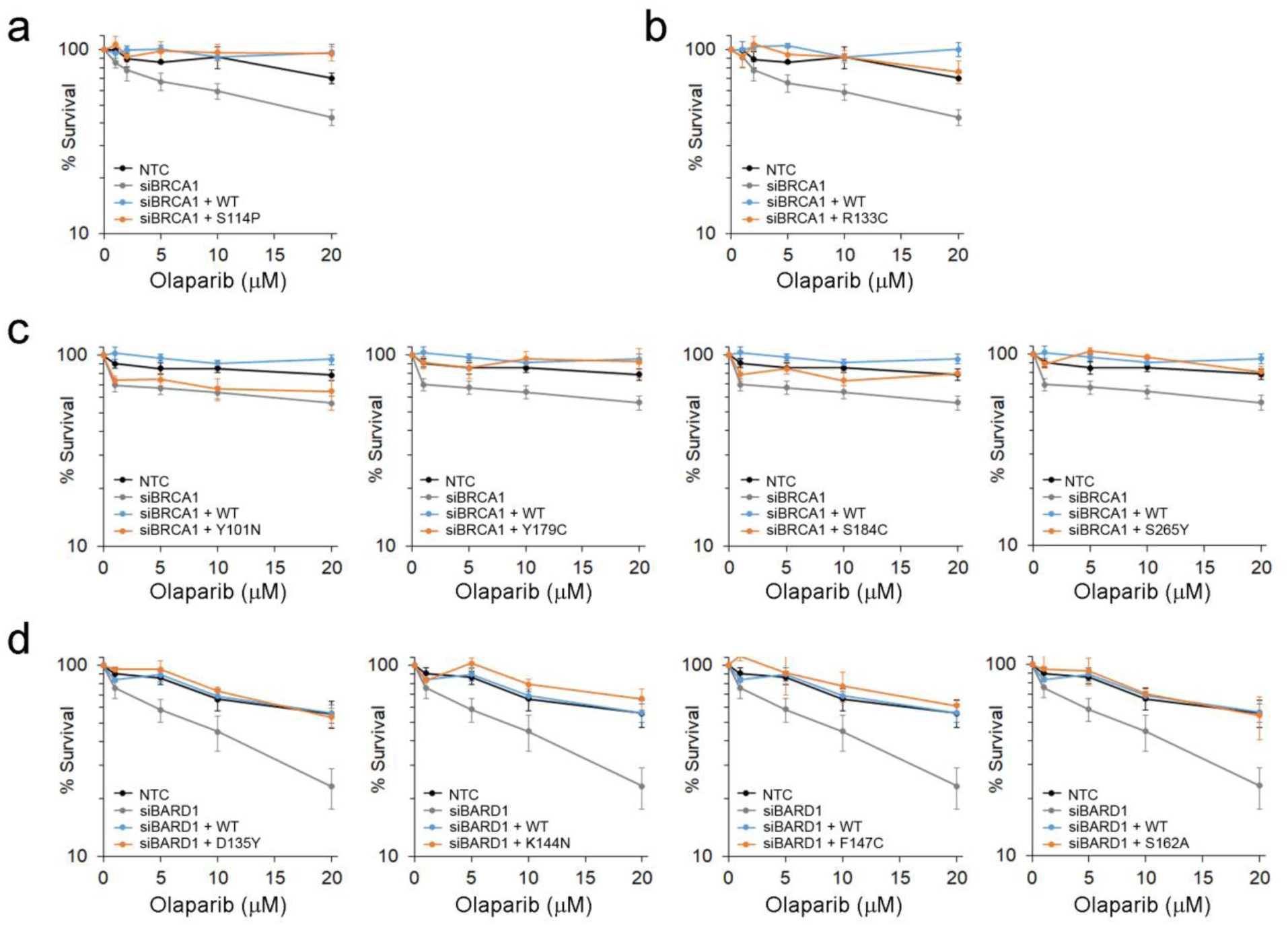
Patient variants define novel functional regions of BRCA1-BARD1 required for fork protection. b. Colony survival following 2 hour treatment with Olaparib was measured in HeLa cells depleted for BRCA1 and complemented with WT or S114P Flag-EGFP-BRCA1. N=4, error bars are SEM. c. Colony survival following 2 hour treatment with Olaparib was measured in HeLa cells depleted for BRCA1 and complemented with WT or R133C Flag-EGFP-BRCA1. N=4, error bars are SEM. d. Colony survival following 2 hour treatment with Olaparib was measured in U20S cells depleted for BRCA1 and complemented with WT or patient variant Flag-EGFP-BRCA1 as indicated. N=3, error bars are SEM. e. Colony survival following 2 hour treatment with Olaparib was measured in U20S cells depleted for BARD1 and complemented with WT or patient variant RFP-Flag-BARD1 as indicated. N=3 for D135Y and S162A. N=4 for K144N and F147C. Error bars are SEM.

**Supplementary table 1.**
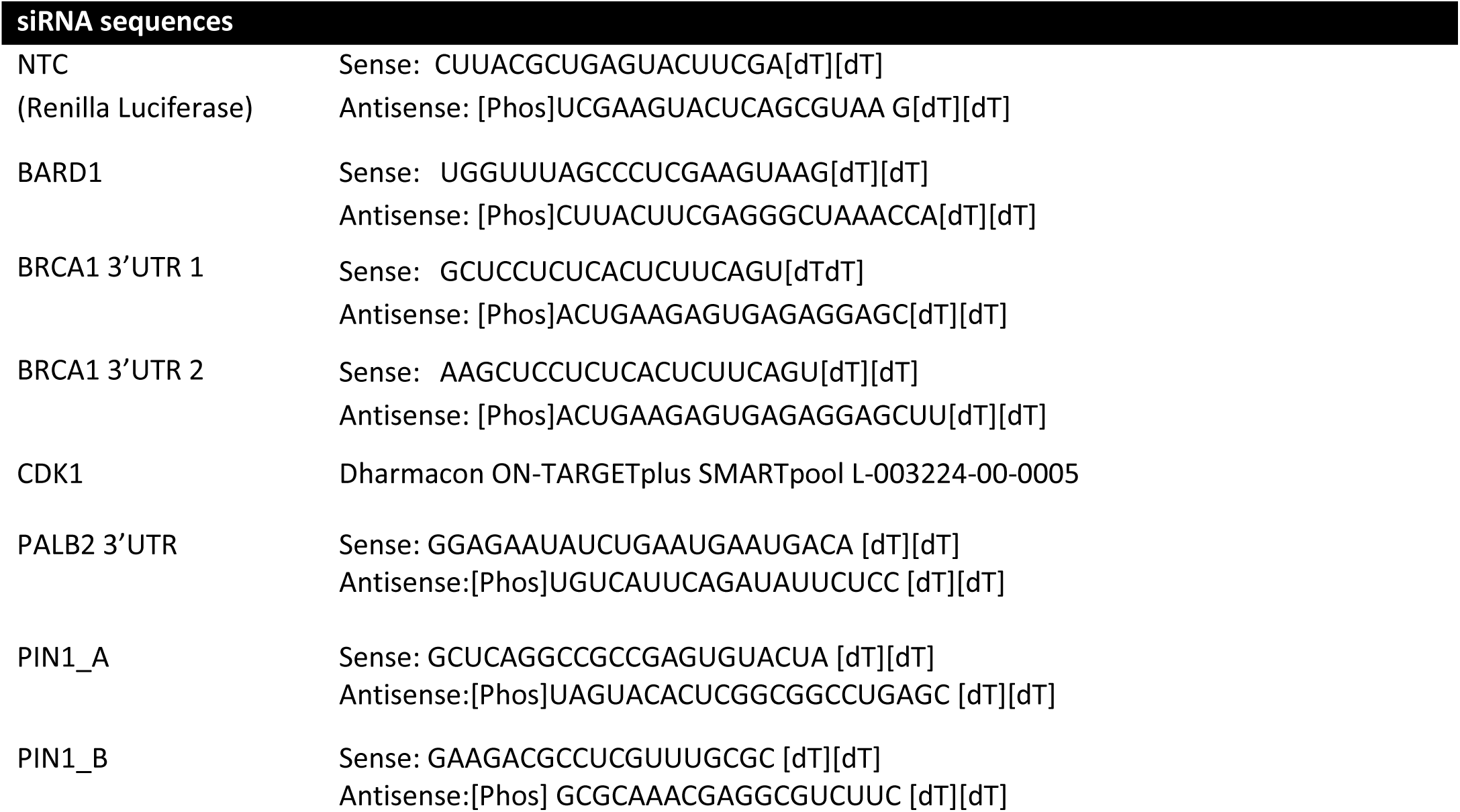
Details of siRNA sequences

**Supplementary table 2.** Details of antibodies and concentrations

| Antibody | Animal | Supplier | Cat. number | Technique | Conc |
| --- | --- | --- | --- | --- | --- |
| β-actin | Rabbit | Abcam | Ab8227 | WB | 1:3000 |
| BRCA1 (D-9) | Mouse | Santa Cruz | Sc6954 | IF | 1:500 |
| BRCA1 (MS110) | Mouse | MERCK Millipore | OP94 | WB | 1:500 |
| pS114-BRCA1 | Rabbit | Genscript | Custom design | WB | 1:1000 |
| CDK1 (A17) | Mouse | Abcam | Ab18 | WB | 1:2000 |
| CldU | Rat | Abcam | Ab6326 | Fibres/IF | 1:2000 |
| Flag (M2) | Mouse | Sigma | F1804 | WB | 1:1000 |
| IdU | Mouse | BD Biosciences | 347580 | Fibres | 1:500 |
| PALB2 | Rabbit | Bethyl | A301.246A | WB | 1:2000 |
| PIN1 | Mouse | R&D Systems | MAB2294 | WB | 1:1000 |
| RAD51 (H-92) | Rabbit | Santa Cruz | SC8349 | WB/IF | 1:1000 |
| RAD51 (Ab-1) | Rabbit | Calbiochem | PC130 | PLA | 1:100 |
| TUBULIN | Mouse | Santa Cruz | sc-5286 | WB | 1:5000 |
| BrdU-Biotin | Mouse | Jackson<br>Immunoresearch | 200-002-211 | PLA | 1:100 |
| BrdU-Biotin | Rabbit | Bethyl | A150-109A | PLA | 1:100 |
| Donkey α Mouse<br>AlexaFluor 488 | Donkey | Life technologies | A21202 | IF | 1:5000 |
| Donkey α Rabbit<br>AlexaFluor 555 | Donkey | Life technologies | A31572 | IF | 1:5000 |
| Donkey α rat<br>AlexaFluor 555 | Donkey | Life technologies | A21434 | IF | 1:5000 |
| Rabbit α Mouse HRP | Rabbit | Dako | P0161 | WB | 1:10000 |
| Swine α Rabbit HRP | Swine | Dako | P0217 | WB | 1:10000 |

**Supplementary Table 3.**
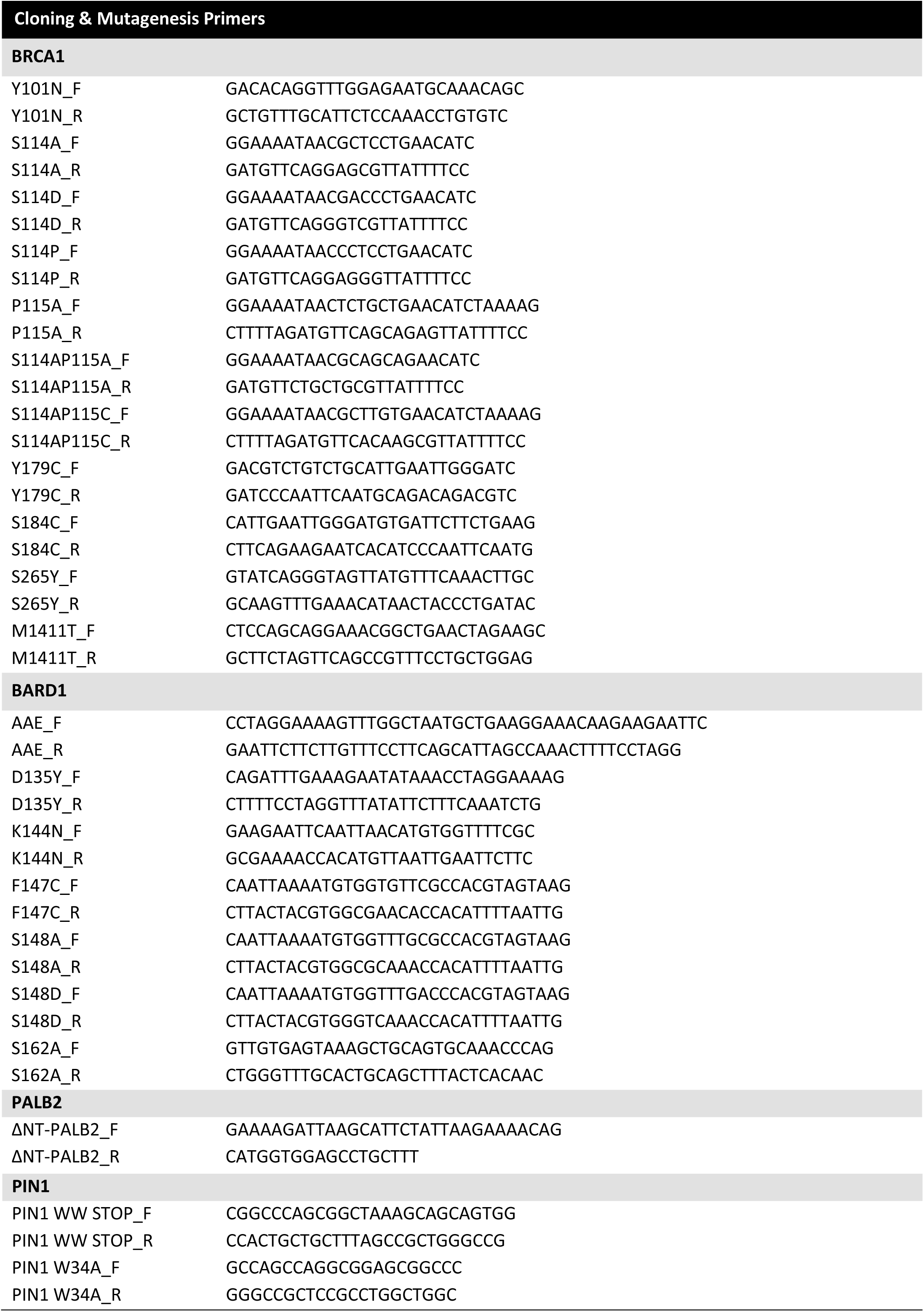
Oligonucleotide primers used for cloning and site-directed mutagenesis

## References

1 Zeman, M. K. & Cimprich, K. A. Causes and consequences of replication stress. Nat Cell Biol 16, 2–9, DOI:10.1038/ncb2897ncb2897 [pii] (2014).

2 Zellweger, R. et al. Rad51-mediated replication fork reversal is a global response to genotoxic treatments in human cells. J Cell Biol 208, 563–579, DOI:10.1083/jcb.201406099 (2015).

3 Mijic, S. et al. Replication fork reversal triggers fork degradation in BRCA2-defective cells. Nat Commun 8, 859, DOI:10.1038/s41467-017-01164-5 (2017).

4 Fumasoni, M., Zwicky, K., Vanoli, F.., Lopes, M. & Branzei, D. Error-free DNA damage tolerance and sister chromatid proximity during DNA replication rely on the Polalpha/Primase/Ctf4 Complex. Mol Cell 57, 812–823, DOI:10.1016/j.molcel.2014.12.038 (2015).

5 Errico, A.., Aze, A. & Costanzo, V. Mta2 promotes Tipin-dependent maintenance of replication fork integrity. Cell Cycle 13, 2120–2128, DOI:10.4161/cc.29157 (2014).

6 Quinet, A.., Lemacon, D. & Vindigni, A. Replication Fork Reversal: Players and Guardians. Mol Cell 68, 830–833, DOI:10.1016/j.molcel.2017.11.022 (2017).

7 Cantor, S. B. & Calvo, J. A. Fork Protection and Therapy Resistance in Hereditary Breast Cancer. Cold Spring Harb Symp Quant Biol, DOI:10.1101/sqb.2017.82.034413 (2018).

8 Schlacher, K. et al. Double-strand break repair-independent role for BRCA2 in blocking stalled replication fork degradation by MRE11. Cell 145, 529–542, DOI:10.1016/j.cell.2011.03.041 (2011).

9 Schlacher, K.., Wu, H. & Jasin, M. A distinct replication fork protection pathway connects Fanconi anemia tumor suppressors to RAD51-BRCA1/2. Cancer Cell 22, 106–116, DOI:10.1016/j.ccr.2012.05.015 (2012).

10 Xu, S. et al. Abro1 maintains genome stability and limits replication stress by protecting replication fork stability. Genes Dev 31, 1469–1482, DOI:10.1101/gad.299172.117 (2017).

11 Leuzzi, G., Marabitti, V.., Pichierri, P. & Franchitto, A. WRNIP1 protects stalled forks from degradation and promotes fork restart after replication stress. Embo J 35, 1437–1451, DOI:10.15252/embj.201593265 (2016).

12 Bhat, K. P. & Cortez, D. RPA and RAD51: fork reversal, fork protection, and genome stability. Nat Struct Mol Biol, DOI:10.1038/s41594-018-0075-z (2018).

13 Higgs, M. R. et al. BOD1L Is Required to Suppress Deleterious Resection of Stressed Replication Forks. Mol Cell 59, 462–477, DOI:10.1016/j.molcel.2015.06.007S1097-2765(15)00446-3 [pii] (2015).

14 Higgs, M. R. et al. Histone Methylation by SETD1A Protects Nascent DNA through the Nucleosome Chaperone Activity of FANCD2. Mol Cell 71, 25–41 e26, DOI:10.1016/j.molcel.2018.05.018 (2018).

15 Hashimoto, Y., Ray Chaudhuri, A.., Lopes, M. & Costanzo, V. Rad51 protects nascent DNA from Mre11-dependent degradation and promotes continuous DNA synthesis. Nat Struct Mol Biol 17, 1305–1311, DOI:10.1038/nsmb.1927 (2010).

16 Wang, A. T. et al. A Dominant Mutation in Human RAD51 Reveals Its Function in DNA Interstrand Crosslink Repair Independent of Homologous Recombination. Mol Cell 59, 478–490, DOI:10.1016/j.molcel.2015.07.009 (2015).

17 Ameziane, N. et al. A novel Fanconi anaemia subtype associated with a dominant-negative mutation in RAD51. Nat Commun 6, 8829, DOI:10.1038/ncomms9829 (2015).

18 Zadorozhny, K. et al. Fanconi-Anemia-Associated Mutations Destabilize RAD51 Filaments and Impair Replication Fork Protection. Cell Rep 21, 333–340, DOI:10.1016/j.celrep.2017.09.062 (2017).

19 Higgs, M. R. & Stewart, G. S. Protection or resection: BOD1L as a novel replication fork protection factor. Nucleus 7, 34–40, DOI:10.1080/19491034.2016.1143183 (2016).

20 Dungrawala, H. et al. RADX Promotes Genome Stability and Modulates Chemosensitivity by Regulating RAD51 at Replication Forks. Mol Cell 67, 374–386 e375, DOI:10.1016/j.molcel.2017.06.023 (2017).

21 Bhat, K. P. & Cortez, D. RPA and RAD51: fork reversal, fork protection, and genome stability. Nat Struct Mol Biol 25, 446–453, DOI:10.1038/s41594-018-0075-z (2018).

22 Ray Chaudhuri, A. et al. Replication fork stability confers chemoresistance in BRCA-deficient cells. Nature 535, 382–387, DOI:10.1038/nature18325 (2016).

23 Yazinski, S. A. et al. ATR inhibition disrupts rewired homologous recombination and fork protection pathways in PARP inhibitor-resistant BRCA-deficient cancer cells. Genes Dev 31, 318–332, DOI:10.1101/gad.290957.116gad.290957.116 [pii](2017).

24 Feng, W. & Jasin, M. BRCA2 suppresses replication stress-induced mitotic and G1 abnormalities through homologous recombination. Nat Commun 8, 525, DOI:10.1038/s41467-017-00634-01 (2017).

25 Dungrawala, H. & Cortez, D. Purification of proteins on newly synthesized DNA using iPOND. Methods Mol Biol 1228, 123–131, DOI:10.1007/978-1-4939-1680-1_10(2015).

26 Sirbu, B. M. et al. Identification of proteins at active, stalled, and collapsed replication forks using isolation of proteins on nascent DNA (iPOND) coupled with mass spectrometry. J Biol Chem 288, 31458–31467, DOI:10.1074/jbc.M113.511337 (2013).

27 Wiest, N. E. & Tomkinson, A. E. Optimization of Native and Formaldehyde iPOND Techniques for Use in Suspension Cells. Methods Enzymol 591, 1–32, DOI:10.1016/bs.mie.2017.03.001 (2017).

28 Pathania, S. et al. BRCA1 haploinsufficiency for replication stress suppression in primary cells. Nat Commun 5, 5496, DOI:10.1038/ncomms6496 (2014).

29 Zhang, F., Fan, Q.., Ren, K. & Andreassen, P. R. PALB2 functionally connects the breast cancer susceptibility proteins BRCA1 and BRCA2. Mol Cancer Res 7, 1110–1118, DOI:1541-7786.MCR-09-0123 [pii] 10.1158/1541-7786.MCR-09-0123 (2009).

30 Zhang, F. et al. PALB2 links BRCA1 and BRCA2 in the DNA-damage response. Curr Biol 19, 524–529, DOI:S0960-9822(09)00723-4 [pii] 10.1016/j.cub.2009.02.018(2009).

31 Sy, S. M., Huen, M. S. & Chen, J. PALB2 is an integral component of the BRCA complex required for homologous recombination repair. Proc Natl Acad Sci U S A 106, 7155–7160, DOI:10.1073/pnas.0811159106 (2009).

32 Zhao, W. et al. BRCA1-BARD1 promotes RAD51-mediated homologous DNA pairing. Nature 550, 360–365, DOI:10.1038/nature24060 (2017).

33 Buisson, R. et al. Breast cancer proteins PALB2 and BRCA2 stimulate polymerase eta in recombination-associated DNA synthesis at blocked replication forks. Cell Rep 6, 553–564, DOI:10.1016/j.celrep.2014.01.009 (2014).

34 Hartford, S. A. et al. Interaction with PALB2 Is Essential for Maintenance of Genomic Integrity by BRCA2. PLoS Genet 12, e1006236, DOI:10.1371/journal.pgen.1006236 (2016).

35 Paull, T. T., Cortez, D., Bowers, B., Elledge, S. J. & Gellert, M. Direct DNA binding by Brca1. Proc Natl Acad Sci U S A 98, 6086–6091 (2001).

36 Densham, R. M. et al. Human BRCA1-BARD1 ubiquitin ligase activity counteracts chromatin barriers to DNA resection. Nat Struct Mol Biol 23, 647–655, DOI:10.1038/nsmb.3236 (2016).

37 Hayami, R. et al. Down-regulation of BRCA1-BARD1 ubiquitin ligase by CDK2. Cancer Res 65, 6–10 (2005).

38 Mertins, P. et al. Proteogenomics connects somatic mutations to signalling in breast cancer. Nature 534, 55–62, DOI:10.1038/nature18003 (2016).

39 Mertins, P. et al. Integrated proteomic analysis of post-translational modifications by serial enrichment. Nat Methods 10, 634–637, DOI:10.1038/nmeth.2518 (2013).

40 Kolinjivadi, A. M. et al. Moonlighting at replication forks - a new life for homologous recombination proteins BRCA1, BRCA2 and RAD51. FEBS Lett 591, 1083–1100, DOI:10.1002/1873-3468.12556 (2017).

41 Steger, M. et al. Prolyl isomerase PIN1 regulates DNA double-strand break repair by counteracting DNA end resection. Molecular Cell 50, 333–343, DOI:10.1016/j.molcel.2013.03.023 (2013).

42 Zheng, H. et al. The prolyl isomerase Pin1 is a regulator of p53 in genotoxic response. Nature 419, 849–853, DOI:10.1038/nature01116nature01116 [pii] (2002).

43 Driver, J. A., Zhou, X. Z. & Lu, K. P. Pin1 dysregulation helps to explain the inverse association between cancer and Alzheimer’s disease. Biochim Biophys Acta 1850, 2069–2076, DOI:10.1016/j.bbagen.2014.12.025 (2015).

44 Dungrawala, H. et al. The Replication Checkpoint Prevents Two Types of Fork Collapse without Regulating Replisome Stability. Mol Cell 59, 998–1010, DOI:10.1016/j.molcel.2015.07.030 (2015).

45 van Drogen, F. et al. Ubiquitylation of cyclin E requires the sequential function of SCF complexes containing distinct hCdc4 isoforms. Mol Cell 23, 37–48, DOI:10.1016/j.molcel.2006.05.020 (2006).

46 Penela, P., Rivas, V.., Salcedo, A. & Mayor, F., Jr. G protein-coupled receptor kinase 2 (GRK2) modulation and cell cycle progression. Proc Natl Acad Sci U S A 107, 1118–1123, DOI:10.1073/pnas.0905778107 (2010).

47 Rustighi, A. et al. The prolyl-isomerase Pin1 is a Notch1 target that enhances Notch1 activation in cancer. Nat Cell Biol 11, 133–142, DOI:ncb1822 [pii] 10.1038/ncb1822 (2009).

48 Liao, Y. et al. Peptidyl-prolyl cis/trans isomerase Pin1 is critical for the regulation of PKB/Akt stability and activation phosphorylation. Oncogene 28, 2436–2445, DOI:onc200998 [pii]10.1038/onc.2009.98 (2009).

49 Weiss, M. S.., Jabs, A. & Hilgenfeld, R. Peptide bonds revisited. Nat Struct Biol 5, 676, DOI:10.1038/1368 (1998).

50 Alderson, T. R., Lee, J. H., Charlier, C.., Ying, J. & Bax, A. Propensity for cis-Proline Formation in Unfolded Proteins. Chembiochem 19, 37–42, DOI:10.1002/cbic.201700548 (2018).

51 Gothel, S. F. & Marahiel, M. A. Peptidyl-prolyl cis-trans isomerases, a superfamily of ubiquitous folding catalysts. Cell Mol Life Sci 55, 423–436, DOI:10.1007/s000180050299 (1999).

52 Ranganathan, R., Lu, K. P.., Hunter, T. & Noel, J. P. Structural and functional analysis of the mitotic rotamase Pin1 suggests substrate recognition is phosphorylation dependent. Cell 89, 875–886 (1997).

53 Pastorino, L. et al. The prolyl isomerase Pin1 regulates amyloid precursor protein processing and amyloid-beta production. Nature 440, 528–534, DOI:nature04543 [pii]10.1038/nature04543 (2006).

54 Lu, K. P. & Zhou, X. Z. The prolyl isomerase PIN1: a pivotal new twist in phosphorylation signalling and disease. Nat Rev Mol Cell Biol 8, 904–916, DOI:nrm2261 [pii]10.1038/nrm2261 (2007).

55 Lu, K. P., Hanes, S. D. & Hunter, T. A human peptidyl-prolyl isomerase essential for regulation of mitosis. Nature 380, 544–547, DOI:10.1038/380544a0(1996).

56 Yaffe, M. B. et al. Sequence-specific and phosphorylation-dependent proline isomerization: a potential mitotic regulatory mechanism. Science 278, 1957–1960 (1997).

57 Lu, P. J., Zhou, X. Z.., Shen, M. & Lu, K. P. Function of WW domains as phosphoserine-or phosphothreonine-binding modules. Science 283, 1325–1328 (1999).

58 Nakamura, K. et al. Proline isomer-specific antibodies reveal the early pathogenic tau conformation in Alzheimer’s disease. Cell 149, 232–244, DOI:10.1016/j.cell.2012.02.016S0092-8674(12)00216-4 [pii] (2012).

59 Hilton, B. A. et al. ATR Plays a Direct Antiapoptotic Role at Mitochondria, which Is Regulated by Prolyl Isomerase Pin1. Mol Cell 60, 35–46, DOI:10.1016/j.molcel.2015.08.008 (2015).

60 Taglialatela, A. et al. Restoration of Replication Fork Stability in BRCA1- and BRCA2-Deficient Cells by Inactivation of SNF2-Family Fork Remodelers. Mol Cell 68, 414–430 e418, DOI:10.1016/j.molcel.2017.09.036 (2017).

61 Petermann, E., Luis Orta, M., Issaeva, N.., Schultz, N. & Helleday, T. Hydroxyurea-Stalled Replication Forks Become Progressively Inactivated and Require Two Different RAD51-Mediated Pathways for Restart and Repair. Molecular Cell 37, 492–502, DOI:10.1016/j.molcel.2010.01.021 (2010).

62 Somyajit, K., Saxena, S., Babu, S.., Mishra, A. & Nagaraju, G. Mammalian RAD51 paralogs protect nascent DNA at stalled forks and mediate replication restart. Nucleic Acids Res 43, 9835–9855, DOI:10.1093/nar/gkv880gkv880 [pii] (2015).

63 Chapman, J. R., Taylor, M. R. & Boulton, S. J. Playing the end game: DNA double-strand break repair pathway choice. Mol Cell 47, 497–510, DOI:10.1016/j.molcel.2012.07.029 (2012).

64 Cerami, E. et al. The cBio cancer genomics portal: an open platform for exploring multidimensional cancer genomics data. Cancer Discov 2, 401–404, DOI:10.1158/2159-8290.CD-12-0095 (2012).

65 Gao, J. et al. Integrative analysis of complex cancer genomics and clinical profiles using the cBioPortal. Sci Signal 6, pl1, DOI:10.1126/scisignal.2004088 (2013).

66 Hedau, S. et al. Novel germline mutations in breast cancer susceptibility genes BRCA1, BRCA2 and p53 gene in breast cancer patients from India. Breast Cancer Res Treat 88, 177–186, DOI:10.1007/s10549-004-0593-8 (2004).

67 Tavtigian, S. V., Byrnes, G. B., Goldgar, D. E. & Thomas, A. Classification of rare missense substitutions, using risk surfaces, with genetic-and molecular-epidemiology applications. Hum Mutat 29, 1342–1354, DOI:10.1002/humu.20896 (2008).

68 Min, S. H. et al. Negative regulation of the stability and tumor suppressor function of Fbw7 by the Pin1 prolyl isomerase. Mol Cell 46, 771–783, DOI:10.1016/j.molcel.2012.04.012 (2012).

69 Yang, W. et al. ERK1/2-dependent phosphorylation and nuclear translocation of PKM2 promotes the Warburg effect. Nat Cell Biol 14, 1295–1304, DOI:10.1038/ncb2629 (2012).

70 Lu, K. P., Finn, G., Lee, T. H. & Nicholson, L. K. Prolyl cis-trans isomerization as a molecular timer. Nat Chem Biol 3, 619–629, DOI:10.1038/nchembio.2007.35 (2007).

71 Esashi, F. et al. CDK-dependent phosphorylation of BRCA2 as a regulatory mechanism for recombinational repair. Nature 434, 598–604, DOI:10.1038/nature03404 (2005).

72 Pefani, D. E. et al. RASSF1A-LATS1 signalling stabilizes replication forks by restricting CDK2-mediated phosphorylation of BRCA2. Nat Cell Biol 16, 962–971, 961-968, DOI:10.1038/ncb3035 (2014).

73 Lemacon, D. et al. MRE11 and EXO1 nucleases degrade reversed forks and elicit MUS81-dependent fork rescue in BRCA2-deficient cells. Nat Commun 8, 860, DOI:10.1038/s41467-017-01180-5 (2017).

74 Ait Saada, A. et al. Unprotected Replication Forks Are Converted into Mitotic Sister Chromatid Bridges. Mol Cell 66, 398–410 e394, DOI:10.1016/j.molcel.2017.04.002 (2017).

75 Tikoo, S. et al. Ubiquitin-dependent recruitment of the Bloom syndrome helicase upon replication stress is required to suppress homologous recombination. EMBO J 32, 1778–1792, DOI:10.1038/emboj.2013.117 (2013).

76 Luijsterburg, M. S. et al. A PALB2-interacting domain in RNF168 couples homologous recombination to DNA break-induced chromatin ubiquitylation. eLife 6, DOI:10.7554/eLife.20922 (2017).

77 Murphy, A. K. et al. Phosphorylated RPA recruits PALB2 to stalled DNA replication forks to facilitate fork recovery. J Cell Biol 206, 493–507, DOI:10.1083/jcb.201404111 (2014).

78 Parplys, A. C. et al. High levels of RAD51 perturb DNA replication elongation and cause unscheduled origin firing due to impaired CHK1 activation. Cell Cycle 14, 3190–3202, DOI:10.1080/15384101.2015.1055996 (2015).

79 Schubert, L. et al. RADX interacts with single-stranded DNA to promote replication fork stability. EMBO Rep 18, 1991–2003, DOI:10.15252/embr.201744877 (2017).

80 Rustighi, A. et al. PIN1 in breast development and cancer: a clinical perspective. Cell Death Differ 24, 200–211, DOI:10.1038/cdd.2016.122 (2017).

81 Bao, L. et al. Prevalent overexpression of prolyl isomerase Pin1 in human cancers. Am J Pathol 164, 1727–1737 (2004).

82 Lu, Z. & Hunter, T. Prolyl isomerase Pin1 in cancer. Cell Res 24, 1033–1049, DOI:10.1038/cr.2014.109 (2014).

83 Liou, Y. C. et al. Loss of Pin1 function in the mouse causes phenotypes resembling cyclin D1-null phenotypes. Proc Natl Acad Sci U S A 99, 1335–1340, DOI:10.1073/pnas.032404099 (2002).

84 Girardini, J. E. et al. A Pin1/mutant p53 axis promotes aggressiveness in breast cancer. Cancer Cell 20, 79–91, DOI:10.1016/j.ccr.2011.06.004 (2011).

85 Ryo, A., Nakamura, M., Wulf, G., Liou, Y. C. & Lu, K. P. Pin1 regulates turnover and subcellular localization of beta-catenin by inhibiting its interaction with APC. Nat Cell Biol 3, 793–801, DOI:10.1038/ncb0901-793 (2001).

86 Farrell, A. S. et al. Pin1 regulates the dynamics of c-Myc DNA binding to facilitate target gene regulation and oncogenesis. Mol Cell Biol 33, 2930–2949, DOI:10.1128/MCB.01455-12 (2013).

87 Rizzolio, F. et al. Retinoblastoma tumor-suppressor protein phosphorylation and inactivation depend on direct interaction with Pin1. Cell Death Differ 19, 1152–1161, DOI:10.1038/cdd.2011.202 (2012).

88 Macheret, M. & Halazonetis, T. D. Intragenic origins due to short G1 phases underlie oncogene-induced DNA replication stress. Nature 555, 112–116, DOI:10.1038/nature25507 (2018).

89 Ahuja, A. K. et al. A short G1 phase imposes constitutive replication stress and fork remodelling in mouse embryonic stem cells. Nat Commun 7, 10660, DOI:10.1038/ncomms10660 (2016).

90 Wulf, G. M. et al. Pin1 is overexpressed in breast cancer and cooperates with Ras signaling in increasing the transcriptional activity of c-Jun towards cyclin D1. Embo J 20, 3459–3472, DOI:10.1093/emboj/20.13.3459 (2001).

91 Wulf, G. M., Liou, Y. C., Ryo, A., Lee, S. W. & Lu, K. P. Role of Pin1 in the regulation of p53 stability and p21 transactivation, and cell cycle checkpoints in response to DNA damage. J Biol Chem 277, 47976–47979, DOI:10.1074/jbc.C200538200C200538200 [pii] (2002).

92 Ryo, A. et al. PIN1 is an E2F target gene essential for Neu/Ras-induced transformation of mammary epithelial cells. Mol Cell Biol 22, 5281–5295 (2002).

93 Zhou, X. Z. & Lu, K. P. The isomerase PIN1 controls numerous cancer-driving pathways and is a unique drug target. Nat Rev Cancer 16, 463–478, DOI:10.1038/nrc.2016.49nrc.2016.49 [pii] (2016).

94 Moore, J. D. & Potter, A. Pin1 inhibitors: Pitfalls, progress and cellular pharmacology. Bioorg Med Chem Lett 23, 4283–4291, DOI:10.1016/j.bmcl.2013.05.088S0960-894X(13)00683-5 [pii] (2013).

95 Wei, S. et al. Active Pin1 is a key target of all-trans retinoic acid in acute promyelocytic leukemia and breast cancer. Nat Med 21, 457–466, DOI:10.1038/nm.3839 (2015).

96 Liao, X. H. et al. Chemical or genetic Pin1 inhibition exerts potent anticancer activity against hepatocellular carcinoma by blocking multiple cancer-driving pathways. Sci Rep 7, 43639, DOI:10.1038/srep43639 (2017).

